# Age-dependent loss of cohesion protection in human oocytes

**DOI:** 10.1101/2023.01.13.523952

**Authors:** Bettina P Mihalas, Gerard H Pieper, Cerys E Currie, David A Kelly, Geraldine M Hartshorne, Andrew D McAinsh, Richard A Anderson, Adele L Marston

## Abstract

Aneuploid human eggs (oocytes) are a major cause of infertility, miscarriage and chromosomal disorders. Such aneuploidies increase greatly as women age, originating from defective linkages between sister-chromatids (cohesion) in meiosis. We found evidence that loss of a specific pool of the cohesin protector protein, shugoshin 2 (Sgo2) contributes to this phenomenon. Our data indicate that Sgo2 preserves sister chromatid cohesion in meiosis by protecting a ‘cohesin bridge’ between sister chromatids. In human oocytes, Sgo2 localizes to both sub-centromere cups and the pericentromeric bridge which spans the sister chromatid junction. Sgo2 normally colocalizes with cohesin, however, in oocytes from older women, Sgo2 is frequently lost specifically from the pericentromeric bridge and sister chromatid cohesion is weakened. Mps1 and Bub1 kinase activities maintain Sgo2 at sub-centromeres and the pericentromeric bridge. Removal of Sgo2 throughout meiosis I by Mps1 inhibition reduces cohesion protection, increasing the incidence of single chromatids at meiosis II. Therefore, Sgo2 deficiency in human oocytes can exacerbate the effects of maternal age by rendering residual cohesin at pericentromeres vulnerable to loss in anaphase I. Our data show that maternal age-dependent loss of Sgo2 at the pericentromere bridge in human oocytes impairs cohesion integrity and contributes to the increased incidence of aneuploidy observed in human oocytes with advanced maternal age.

## Introduction

In humans, infertility and miscarriage are common, and a leading cause is aneuploidy in the egg (oocyte)^1^. Even among women at peak fertile age, over 20% of oocytes are aneuploid, while at advanced maternal age (>32 years), aneuploidy affects more than 50% of oocytes^2^. This aneuploidy arises predominantly from errors in meiosis, the specialized cell division that generates oocytes which have half the number of chromosomes of the parental cell. Meiosis I segregates the homologous chromosomes, while meiosis II segregates the sister chromatids^3^. Accurate sequential execution of meiosis I and II requires the establishment of cohesin complexes during DNA replication to link sister chromatids together and the preservation of this cohesin until chromosomes are ready to segregate. In mammalian oocytes, cohesin is established *in utero* after which oocytes enter a long arrest in meiotic prophase I: meiosis is not resumed until the oocyte is ovulated, potentially several decades later. Evidence from mice indicates that there is no cohesin turnover during this period^4,5^, suggesting that cohesin laid down in the fetus must last throughout the female reproductive lifespan, up to age ~50 years in humans. However, female fertility declines before this, largely due to the loss of oocyte euploidy. Upon resumption of meiosis, cohesin is lost in two steps. During meiosis I, loss of chromosomal arm cohesin triggers homolog segregation. However, centromeric cohesin is protected during meiosis I and lost only in meiosis II to allow sister chromatid segregation. Reduced sister-chromatid cohesion is a key driver of aneuploidy in human oocytes, particularly with increased maternal age^2,6–9^. Therefore, the molecular origin of this age-dependent cohesin loss in human oocytes requires investigation.

In most organisms, a meiosis-specific variant of cohesin containing the Rec8 kleisin confers sister chromatid cohesion and Rec8 cleavage by separase destroys cohesion in two steps^10,11^. In meiosis I, phosphorylation-dependent cleavage of Rec8 by separase occurs only on chromosome arms, triggering homologous chromosome segregation^12–18^. However, pericentromeric Rec8 is protected from phosphorylation, and therefore cleavage, by shugoshin family proteins which recruit protein phosphatase 2A (PP2A) to the pericentromeres^19–21^. Retention of pericentromeric cohesion in meiosis I permits sister chromatid alignment in meiosis II followed by separase re-activation, cohesin de-protection and sister chromatid segregation^22–24^. In mouse oocytes, shugoshin 2 (Sgo2) protects Rec8 from separase: *sgo2* knockout mice are viable but infertile because all cohesin is cleaved in anaphase I, resulting in random segregation of sister chromatids at meiosis II^21,25,26^.

Consistent with its role in pericentromeric cohesin protection, shugoshin localizes to proximal to centromeres and spanning the junction between sister centromeres^27^. This sister chromatid junction corresponds to the so-called inner centromere where both pericentromeric heterochromatin and protected cohesin reside. The pericentromeric localization of shugoshin in both meiosis and mitosis is under control of the Bub1 and Mps1 kinases (reviewed in ^28^). In cultured human mitotic cells, Mps1 phosphorylates the kinetochore protein Knl1 to recruit Bub1^29,30^. Bub1 in turn phosphorylates histone H2A on Thr120 to provide a docking site for Sgo1 and thereby protect mitotic cohesin from release via the separase-independent prophase pathway^31,32^. In contrast, in mouse oocytes, although Bub1 kinase activity promotes Sgo2 localization at centromeres, it appears dispensable for fertility^27,33^. Instead, cohesin protection in mouse oocytes is achieved through Mps1-dependent Sgo2 localization^27^. Cohesin-deficient mouse oocytes show aneuploidies similar to the age-related decline in human oocytes^34^. Reduced levels of chromosomal cohesin and Sgo2 with increased maternal age^35–37^ have been observed in some mouse strains, however female mice do not show reproductive aging to the same extent as humans. Therefore, understanding the role of cohesin protection in counteracting age-related aneuploidy requires direct analysis of human oocytes. The discovery of a frameshift mutation in *SGO2* as the likely cause of ovarian insufficiency and infertility in a female patient^38^ imply a critical role for Sgo2 in human oogenesis, but the effects of aging on Sgo2 and cohesin protection have not been analysed.

Here we use high resolution microscopy to show that in human metaphase I and II oocytes, the two sister kinetochores are embedded within Sgo2 cups which connect through a pericentromeric bridge spanning the sister chromatid junction. Oocytes from older women frequently show a specific reduction of Sgo2 at the bridge. Furthermore, inhibition of Mps1 in oocytes from younger women causes removal of Sgo2 from the pericentromeric bridge and loss of cohesion protection, recapitulating the phenotype of older oocytes. These findings support a model where age-dependent decline in association of Sgo2 with the pericentromere bridge makes pericentromeric cohesion vulnerable to premature loss, a major cause of age-related aneuploidy in humans. It follows that methods preserving/supplementing Sgo2 may offer potential to support fertility while reducing oocyte aneuploidy in women of older reproductive age.

## Results

### Sgo2 cups and bridges sister kinetochores in human oocytes

To understand whether Sgo2 could protect cohesin during the long prophase I arrest and meiotic divisions, we examined its localization in whole human oocytes at different meiotic stages (Figure S1A). Sgo2 signal was detected in the germinal vesicle nucleus already during prophase I but became enriched on chromosomes only following germinal vesicle breakdown (GVBD), marking the end of prophase I arrest. By prometaphase I, Sgo2 was focused around the inner kinetochore marker, CENP-C. Sgo2 was also localized near the kinetochore marker in metaphase I and also in metaphase II, at which point oocytes naturally arrest awaiting fertilisation.

To observe Sgo2 localization in more detail, we used high resolution microscopy on whole human oocytes at metaphase I to obtain 3D reconstructions and analysed individual bivalents (homologous chromosomes connected by chiasmata). Human Sgo2 surrounded centromeres and spanned the junction between sister centromeres, which we refer to as the pericentromeric bridge (Figure 1A and B; age 30 years). Centromeric Sgo2 surrounds, but does not overlap, the two sister centromeres/inner kinetochores (as identified by CREST or CenpC staining), forming cup-like structures that are frequently connected by a Sgo2 bridge (Figure 1B). In metaphase II, where sister kinetochores form a back-to-back configuration, the Sgo2 bridge was found to span the distance between the Sgo2 centromeric cups (Figure 1C and D; age 29 years). Line scans of metaphase II chromosomes further suggested the existence of separate centromeric and bridge Sgo2 pools (Figure S1B). Therefore, human Sgo2 is positioned at sub-centromere cups and a pericentromeric chromatin bridge during the oocyte meiotic divisions, consistent with a role in pericentromeric cohesin protection.

**Figure 1.**
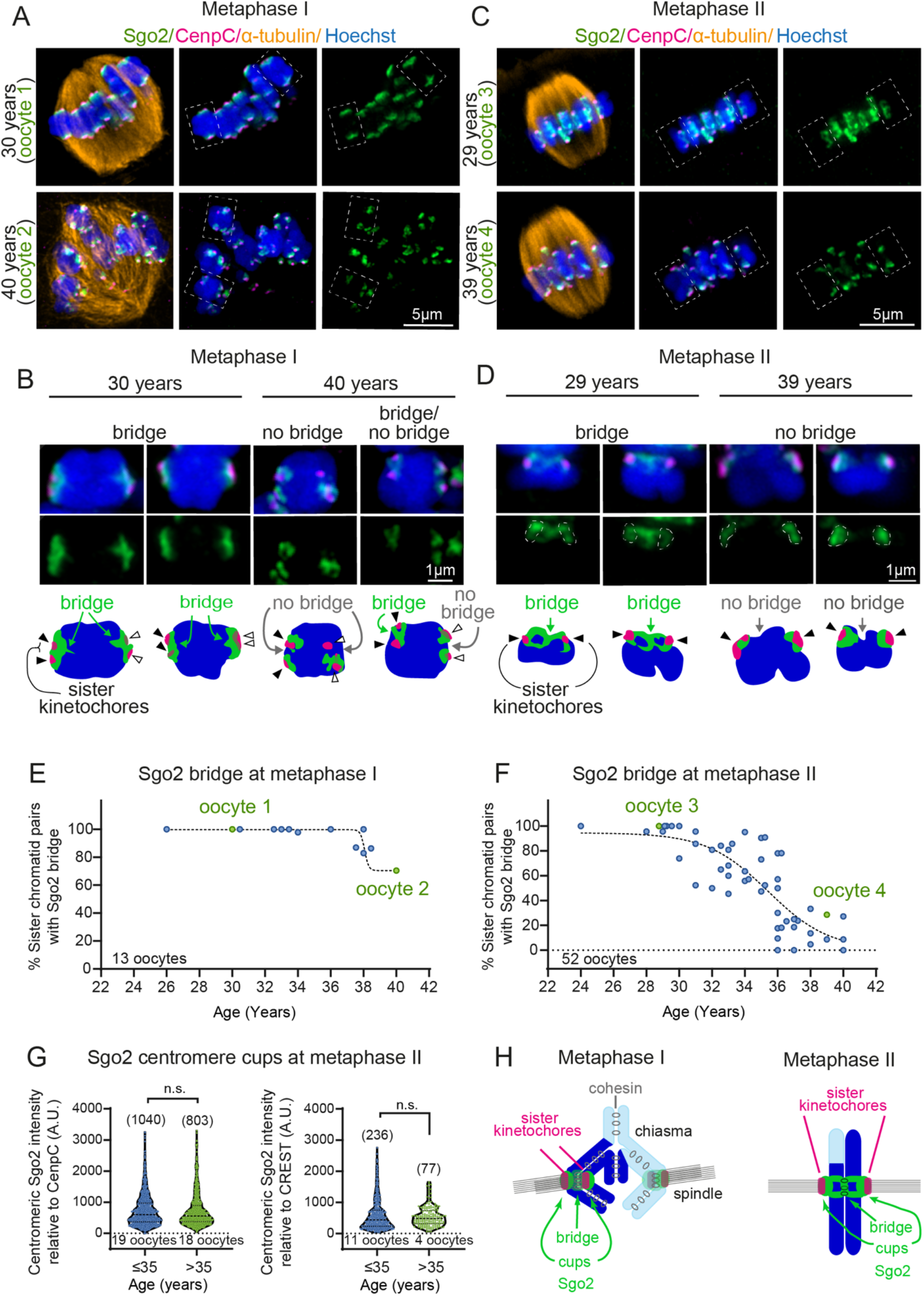
Age-dependent loss of human Sgo2 from the pericentromeric bridge, but not centromeres, in meiosis I oocyte and meiosis II oocytes. (A-D) Human Sgo2 localizes to centromeric cups and an inter-sister pericentromere bridge in metaphase I oocytes and metaphase II oocytes of younger women, but is lost from the bridge with age. Representative images of human metaphase I (A and B) and metaphase II oocytes (C and D) from younger (age 30 and 29 years) and older women (age 40 and 39 years). Sgo2 (green), inner kinetochores (CenpC, magenta), microtubules (a-tubulin, orange) and chromosomes (Hoeschst, blue) are shown. White dashed line boxes indicate chromosomes further magnified in (B) metaphase I and (D) metaphase II. Centromeric localization of Sgo2 is bounded by dashed lines. Interpretations of localizations observed are indicated in the schematics below. (E and F) The percentage of chromatids per oocyte with Sgo2 localization at the pericentromeric bridge was scored relative to the woman’s age at MI (E) and MII (F). Data were fit to a Sigmodal, 4PL curve (metaphase I; R^2^ = 0.84, metaphase II; R^2^ = 0.67). Oocytes shown in (A-D) are labelled (green circles). (G) Centromeric Sgo2 signal is not significantly altered with age. The relative intensity of the centromeric pool of Sgo2 in metaphase II oocytes from younger (age ≤ 35 years) or older women (age >35 years) was measured in arbitrary units (A.U.) relative to the inner kinetochore (CENP-C) or centromere (CREST) markers. Plots show median (dashed black line), 25th and 75th percentiles (dotted black lines). *P* values were calculated using the Mann-Whitney test: CenpC (*P* = 0.37) and CREST (*P* = 0.15). n.s. = not significant. (H) Schematic summarises Sgo2 localization in younger women in metaphase I and II oocytes.

### Age-dependent loss of Sgo2 from the pericentromeric bridge

Although Sgo2 centromeric cups were consistently observed, a Sgo2 bridge was not always present (Figure 1A-D; age 40 and 39 years), and line scans confirmed the specific absence of the Sgo2 pool at the pericentromeric bridge on some chromosomes (Figure S1B). Scoring the frequency of sister chromatid pairs with an intact Sgo2 bridge in meiosis I (Figure 1E) and II (Figure 1F) revealed a clear relationship between the presence of the Sgo2 bridge and the woman’s age. In oocytes from women aged 30 years and under, a Sgo2 bridge was observed at both metaphase I and metaphase II on almost all sister chromatid pairs (Figure 1E and F). At higher maternal ages, however, the frequency of sister chromatids with a Sgo2 bridge was decreased. This effect was most obvious in meiosis II oocytes, where the back-to-back configuration of sister kinetochores facilitates observation of the pericentromeric region. Accordingly, at over 30 years of age the frequency of chromosomes lacking a Sgo2 bridge in meiosis II increased and oocytes from most women aged 36 years or over lacked an Sgo2 bridge on the majority of sister chromatid pairs (Figure 1F). An increased frequency of chromosomes lacking the Sgo2 bridge was also observed in meiosis I oocytes from women over 35 years of age, despite the challenges of distinguishing centromeric and bridge Sgo2 when kinetochores are in the side-by-side orientation (Figure 1E). The age-dependence of the Sgo2 bridge was also not affected by whether the oocytes had been acquired during *in vitro* fertilisation (IVF) or intracytoplasmic sperm injection (ICSI) treatment (Figure S1C). In contrast, measurement of centromeric Sgo2 signal (normalised to CREST or CenpC) on meiosis II chromosomes revealed no significant difference with age (Figure 1G). Therefore, aging leads to a specific decline in the Sgo2 pool at the pericentromeric bridge. We conclude that in human oocytes at metaphase I and II, Sgo2 exists in two chromosomal pools, the cups and the bridge, and that bridge Sgo2 is particularly vulnerable to aging (Figure 1H).

### Absence of the Sgo2 bridge is associated with increased inter-sister kinetochore distance

Increased inter-sister kinetochore distance, indicative of a loss of centromeric cohesion, has been observed in aged human oocytes^7–9^. To understand the relationship between cohesion loss and the lack of a Sgo2 bridge, we measured the inter-sister kinetochore distance at metaphase I and II and correlated this to the presence or absence of Sgo2 staining at the bridge (Figure 2A-F; Note that in metaphase I, we defined the bridge as continuous Sgo2 staining between the CenpC/Crest centromere signals). As expected, we observed a clear correlation between sister kinetochore distance and female age at both metaphase I and metaphase II (Figure 2B and E), which was consistent whether the women had undergone IVF or ICSI treatment (Figure S2), confirming that centromeric cohesion weakens with age. Additionally, meiosis I and II sister kinetochores that lack a Sgo2 bridge tended to be further apart (Figure 2B and E). Although sister kinetochore pairs with a Sgo2 bridge showed increased separation in meiosis I oocytes from older (age >35 years) compared to younger (age ≤35 years) women, the greatest inter-sister kinetochore distance was observed on those pairs lacking a Sgo2 bridge (Figure 2C). Similarly, even in metaphase II oocytes from young (age ≤35 years) women, sister chromatid pairs lacking a Sgo2 bridge were further apart than pairs with a Sgo2 bridge (Figure 2F). In contrast, centromeric Sgo2 signal did not correlate with inter-sister kinetochore distance (Figure 2G). Therefore, regardless of oocyte age, decreased centromeric cohesion is associated with specific loss of the bridge, but not centromeric, Sgo2.

**Figure 2.**
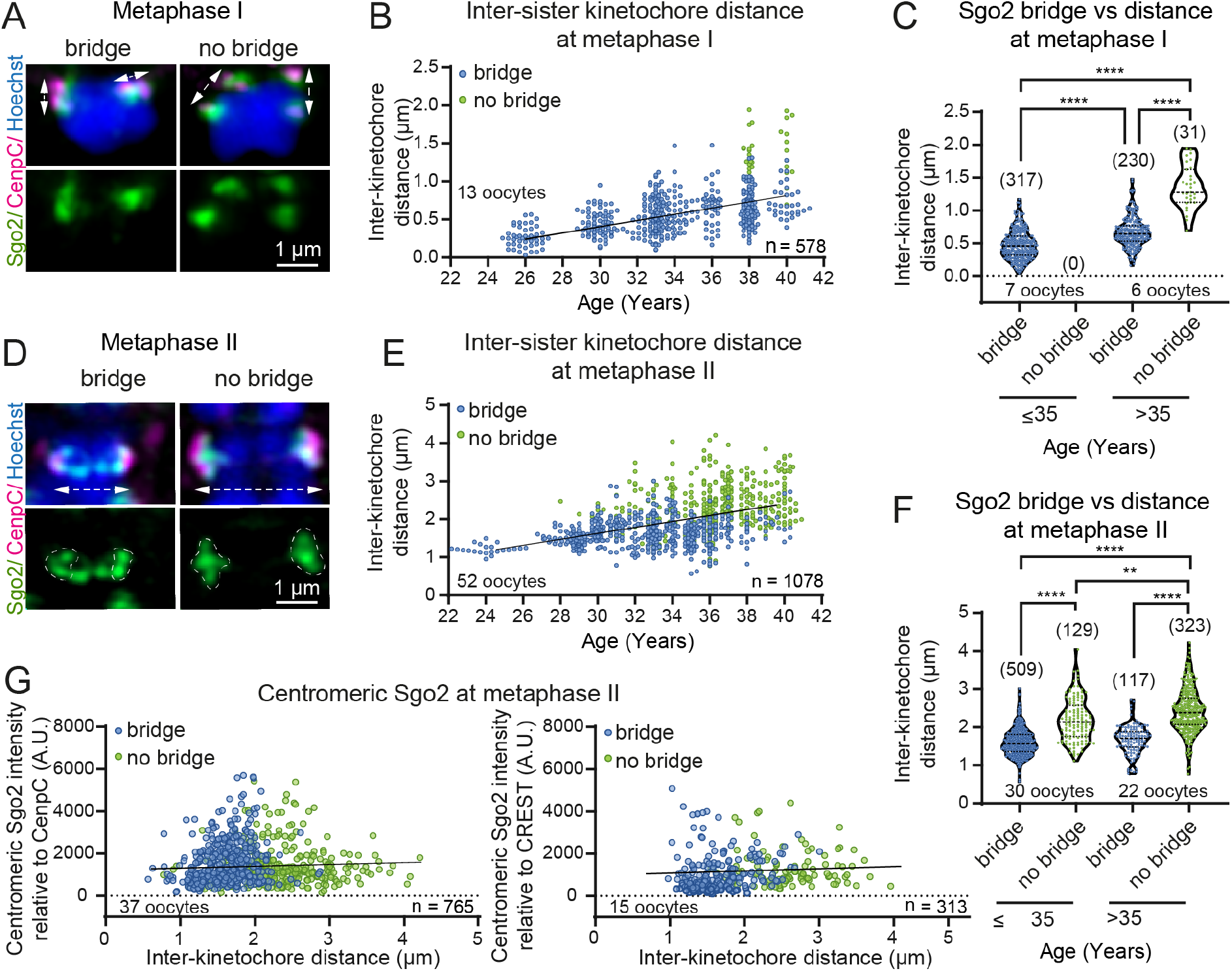
Loss of Sgo2 at the pericentromeric bridge with increased inter-sister kinetochore distance. (A-F) Loss of Sgo2 at the pericentromeric bridge is associated with increased inter-sister kinetochore distance in metaphase I and metaphase II. (A) Inter-sister kinetochore distance (white dashed arrows) was determined on metaphase I chromosomes and related to the presence of the Sgo2 bridge. (B) Increase in inter-sister kinetochore distance at metaphase I with female age for chromosomes with a Sgo2 bridge (blue) or no bridge (green). Data was fit to a linear regression (R^2^ = 0.81; *P* <0.0001). (C) Inter-sister kinetochore distance for sister chromatid pairs with a Sgo2 bridge or no bridge at metaphase I in younger (age ≤ 35 years) or older women (age >35 years). Plots show median (dashed black line), 25th and 75th percentiles (dotted black lines). Statistical analyses were performed using the Kruskal-Wallis test (\*\*\*\**P* <0.0001). (D) Inter-sister kinetochore distance (white dashed arrows) was determined on metaphase II chromosomes and related to the presence of the Sgo2 bridge. (E) Increase in inter-sister kinetochore distance at metaphase II with female age for sister chromatid pairs with a Sgo2 bridge (blue) or no bridge (green). Data was fit to a linear regression (R^2^ = 0.47; *P* <0.0001). (F) Inter-sister kinetochore distance where the Sgo2 bridge is present or absent at metaphase II in younger (age ≤ 35 years) or older women (age >35 years). Statistical analyses were performed using the Kruskal-Wallis test (\*\*\*\**P* <0.0001, \*\**P* = 0.0017). Plots show median (dashed black line), 25th and 75th percentiles (dotted black lines). (G) Centromeric Sgo2 does not correlate with inter-sister kinetochore distance. The relative intensity of the centromeric pool of Sgo2 metaphase II oocytes was measured in arbitrary units (A.U.) relative to the kinetochore markers CenpC (*P* =0.23; R^2^ = 0.0019) and CREST (*P* = 0.052; R^2^ = 0.012). Data was fit to a linear regression.

### Incidence of single chromatids in meiosis II oocytes is higher where many sister chromatid pairs lack a Sgo2 bridge

In the most extreme, defective centromeric cohesion would abolish all linkages between sister chromatids resulting in the presence of single chromatids in naturally arrested metaphase II oocytes. To further understand the consequences of defective sister chromatid cohesion in aged human oocytes (Figure 2E), we scored the number of single chromatids in our 3D metaphase II reconstructions (Figure 3A). As expected^7^, this revealed a sharp increase in single chromatids within oocytes from women >35 years of age, independent of their treatment (Figure 3B; Figure S3A). More single chromatids were observed in oocytes with an elevated frequency of sister chromatid pairs lacking a Sgo2 bridge (Figure 3C). In contrast, centromeric Sgo2 signal was comparable on single and paired sister chromatids (Figure 3D), indicating that association of Sgo2 with centromeric cups is independent of cohesion. Single chromatids were rarely detected in meiosis I oocytes from women of any age (Figure S3B). This indicates that chromosome arms have sufficient cohesion to ensure chromatids remain juxtaposed and that it is only after loss of arm cohesion, when sister chromatids rely solely on centromeric cohesion, that the consequences of defective centromeric cohesion are apparent.

**Figure 3.**
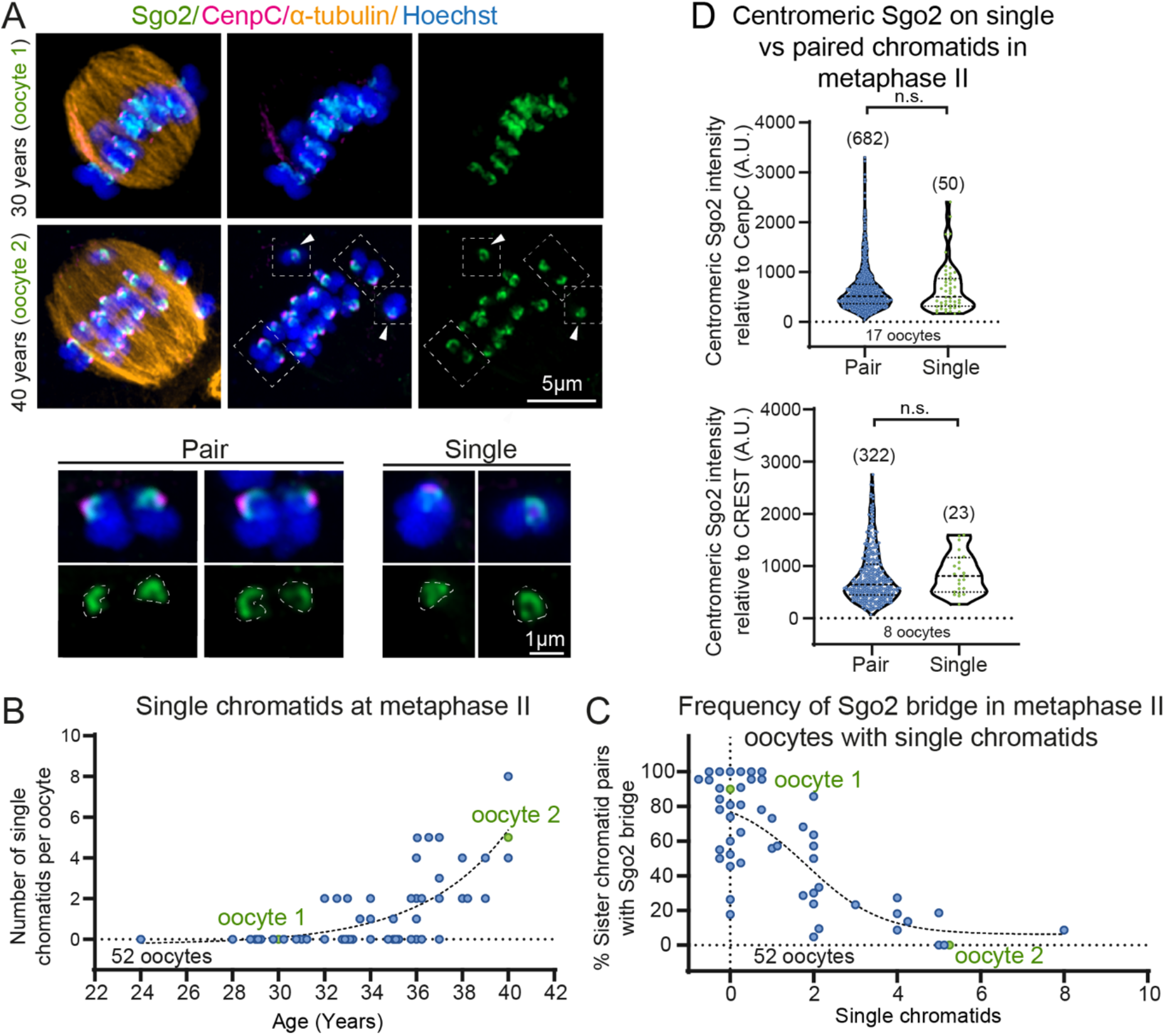
Loss of cohesion in metaphase II oocytes with age and absence of bridge Sgo2. (A) Representative image showing presence of single chromatids (arrows) in human metaphase II oocytes from an older (age 40) women, compared to a younger (age 30) woman without single chromatids. White boxes with dashed lines indicate examples of paired and single chromosomes in oocyte 2. Sgo2 (green), inner kinetochores (CenpC; magenta), microtubules (a-tubulin; orange) and chromosomes (Hoechst; blue) are shown. (B) Increase of single chromatids in metaphase II oocytes with maternal age. The number of single chromatids was scored relative to woman’s age. Data were fit to a Sigmodal, 4PL curve (R^2^ = 0.57). Oocytes used in representative images are labelled in the graphs. (C) Increased number of single chromatids in metaphase II oocytes with an increased fraction of sister chromatid pairs lacking a Sgo2 bridge. Data were fit to a Sigmodal, 4PL curve (R^2^ = 0.60). (D) The relative intensity of the centromeric pool of Sgo2 between paired and single chromatids was measured only from oocytes that had single chromatids. Sgo2 intensity was measured in arbitrary units (A.U.) relative to the kinetochore markers CenpC (*P* = 0.96, Mann-Whitney test) and CREST (*P* = 0.24, Mann-Whitney test). Plots show median (dashed black line), 25th and 75th percentiles (dotted black lines). *P* values were calculated using the Mann-Whitney test. n.s., Not significant.

### Sgo2 co-localizes with PP2A and Rec8 at pericentromeres in meiosis II

In yeast and mouse, it is well-established that shugoshins protect Rec8-cohesin until anaphase II of meiosis through recruitment of PP2A to pericentromeric regions^19–21,39^. Our findings above revealed a relationship between centromeric cohesion and the Sgo2 bridge, suggesting that this pool of Sgo2 might be most relevant in Rec8 protection. To test this idea further, we compared Sgo2 and Rec8 localization on spread metaphase II chromosomes from human oocytes. At this stage, we expected arm cohesin to have been lost so only centromeric cohesin remains. Consistently, Rec8 was detected only in pericentromeric regions (Figure 4A). As in whole oocytes, Sgo2 encircled centromeres (as identified by CenpC staining) and frequently formed a pericentromeric bridge across the sister centromere junction (Figure 4A). Importantly, the Sgo2 signal partially overlapped with Rec8 and there was a strong correlation between the presence of a Sgo2 bridge and Rec8 across the sister chromatid junction (Figure 4B). Similarly, PP2A co-localized with Sgo2 at the pericentromere bridge on metaphase II chromosome spreads, but was absent from the pericentromere on chromosomes that lacked a Sgo2 bridge (Figure 4C and D). These findings reveal co-localization of Rec8-cohesin, Sgo2 and PP2A at the pericentromeric bridge spanning the sister chromatid junction in human meiosis II oocytes and further implicate the Sgo2 pool at the bridge in protecting pericentromeric cohesion.

**Figure 4.**
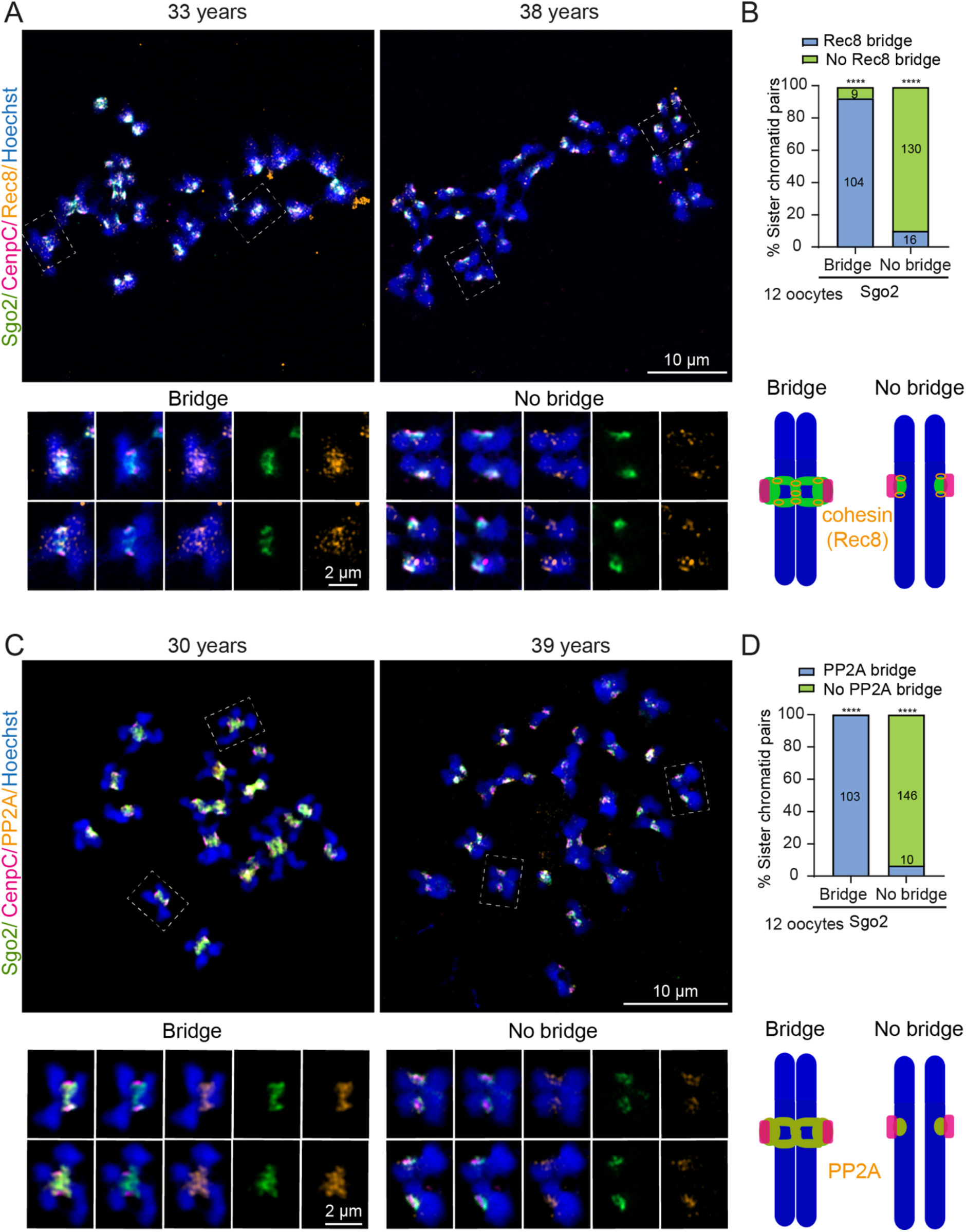
Co-localization of PP2A and cohesin with the Sgo2 pericentromere bridge on human metaphase II chromosomes. (A and B) Chromosome spreads of metaphase II-arrested human oocytes were stained with antibodies against Sgo2 (green), inner kinetochores (CenpC, magenta) and cohesin (Rec8, orange). Chromosomes were stained with Hoechst (blue). (A) Representative images with white dashed line boxes indicating representative chromosome figures shown at higher magnification on the right. (B) Localization of Rec8 in the pericentromere bridge was scored for sister chromatid pairs with and without a Sgo2 bridge. Schematic representations of the data are shown below. (C and D) Chromosome spreads of metaphase II-arrested human oocytes were stained with antibodies against Sgo2 (green), inner kinetochores (CenpC, magenta) and PP2A (orange). Chromosomes were stained with Hoechst (blue). (C) Representative images with white dashed line boxes indicating representative chromosome figures shown at higher magnification on the right. (D) Localization of PP2A in the pericentromere bridge was scored for sister chromatid pairs with and without a Sgo2 bridge. Schematic representations of the data are shown below.

### Sgo2 localization requires Mps1 and Bub1 activity

In human mitotic cells and mouse oocytes, Mps1 and Bub1 kinase activities promote the localization of shugoshin proteins to pericentromeres^27,31,32,40–42^. To understand the role of these kinases in localizing human Sgo2 to the centromere and bridge in oocytes, we used specific inhibitors. Human metaphase II oocytes were incubated with the Mps1 inhibitor, Reversine^43^ for 16 h before fixing and staining by immunofluorescence (Figure 5A and B). Mps1 inhibition in metaphase II oocytes resulted in loss of the Sgo2 bridge, together with a significant reduction in centromeric Sgo2 signal (Figure 5B-D). Mps1 is required for chromosome biorientation and error correction in mitotic cells and its inhibition is predicted to increase attachment of sister kinetochores to microtubules from the same pole^29^. Accordingly, we observed a reduction in inter-kinetochore distance upon Mps1 inhibition in metaphase II cells, while the number of single chromatids was not increased (Figure S4). Mps1 activity localizes Bub1 to kinetochores in human mitotic cells^44,45^ and oocytes^7^.

**Figure 5.**
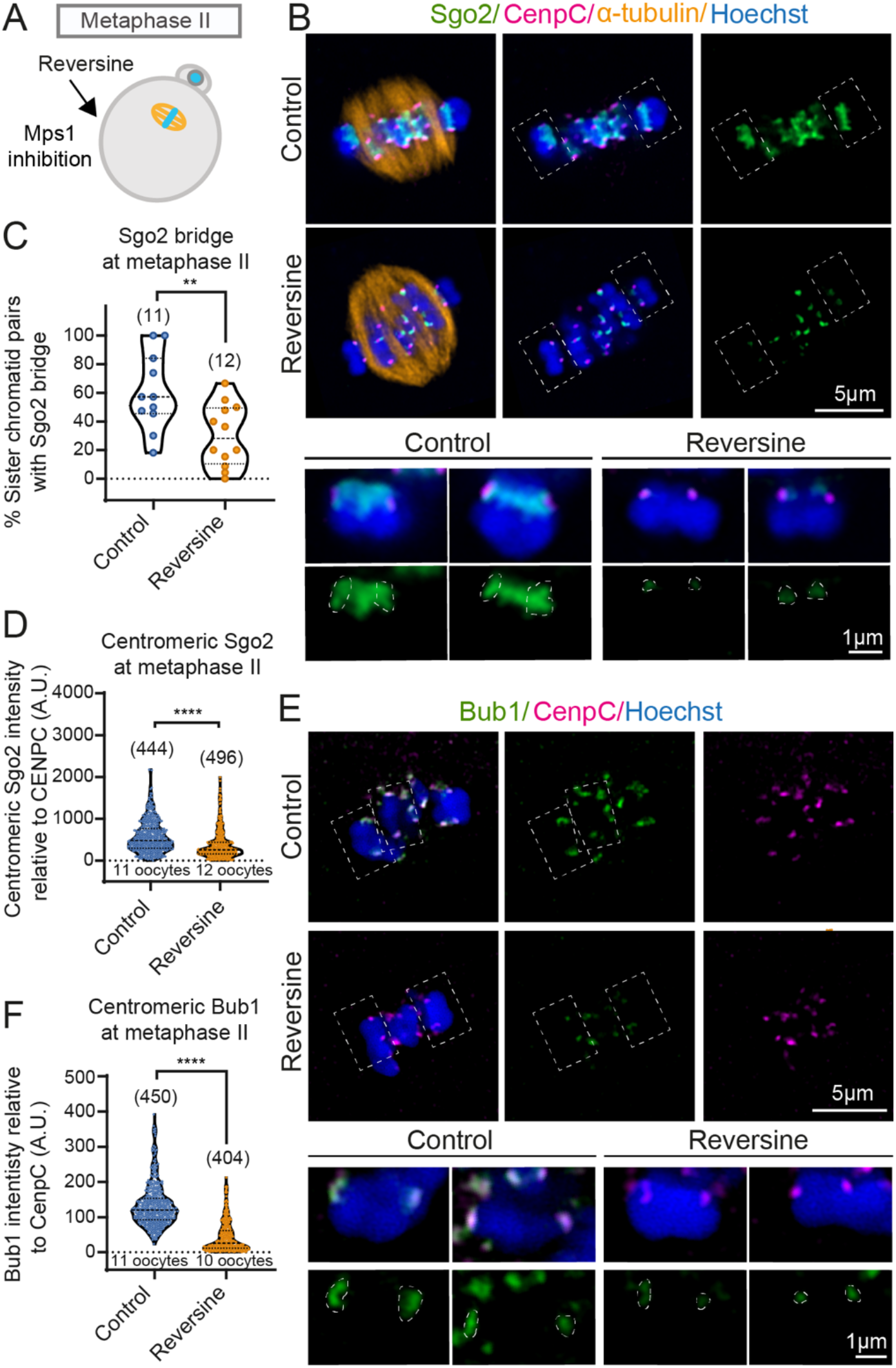
Sgo2 localization to the bridge and centromere requires MPS1 activity. (A-D) Inhibition of Mps1 in metaphase II oocytes impairs Sgo2 localization. (A) Scheme of the experiment. Metaphase II eggs from women aged ≤ 36 years were treated with 500nM Reversine^43^ (to inhibit Mps1) or DMSO (control) overnight and fixed. (B) Representative images of control and reversine-treated metaphase II oocytes after immunostaining with antibodies against Sgo2 (green), CenpC (inner kinetochores, magenta) and a-tubulin (microtubules, orange). White boxes with dashed lines indicate chromosomes that have been further magnified below. (C) The percentage of chromatids per oocyte with Sgo2 localization at the pericentromeric bridge was scored in control and reversine-treated metaphase II oocytes (\*\**P* =0.0084; Welch’s t test). (D) The relative intensity of the centromeric pool of Sgo2 in metaphase II oocytes relative to CenpC in control and reversine-treated oocytes (\*\*\*\**P* <0.0001; Mann-Whitney test). (E and F) Mps1 activity is required for Bub1 localization to kinetochores. Metaphase II oocytes treated with reversine as in (A) were immunostained with antibodies against Bub1 (green), CenpC (inner kinetochore; magenta), and counter stained with Hoechst (blue) to visualise chromosomes. (E) Representative images of Bub1 localisation in control and reversine-treated metaphase II oocytes. White boxes with dashed lines represent chromosomes that have been further magnified below. White dashed circles show examples of area selections for Bub1 intensity measurements. (F) The relative intensity of the centromeric Bub1 in metaphase II oocytes from control and reversine-treated oocytes relative to CenpC (\*\*\*\**P* <0.0001; Mann-Whitney test). Plots show median (dashed black line), 25th and 75th percentiles (dotted black lines).

Consistent with this, we found that the intensity of centromere-proximal Bub1 signal was greatly reduced after treating metaphase II oocytes with Reversine (Figure 5E and F). We therefore tested the role of Bub1 in Sgo2 localization by treatment with the Bub1 inhibitor Bay-320^46^ (Figure 6A). Bub1 inhibition in metaphase II oocytes resulted in almost complete loss of the Sgo2 signal at both the centromeric cups and bridge (Figure 6B-D), though this was not accompanied by an increase in either inter-sister kinetochore distance or the frequency of single chromatids. We conclude that the activities of both Mps1 and Bub1 contribute to localising Sgo2 at both the centromeric cups and inter-sister kinetochore bridge in human metaphase II oocytes and that the role of Mps1 may be executed through Bub1 recruitment (Figure 6E).

**Figure 6.**
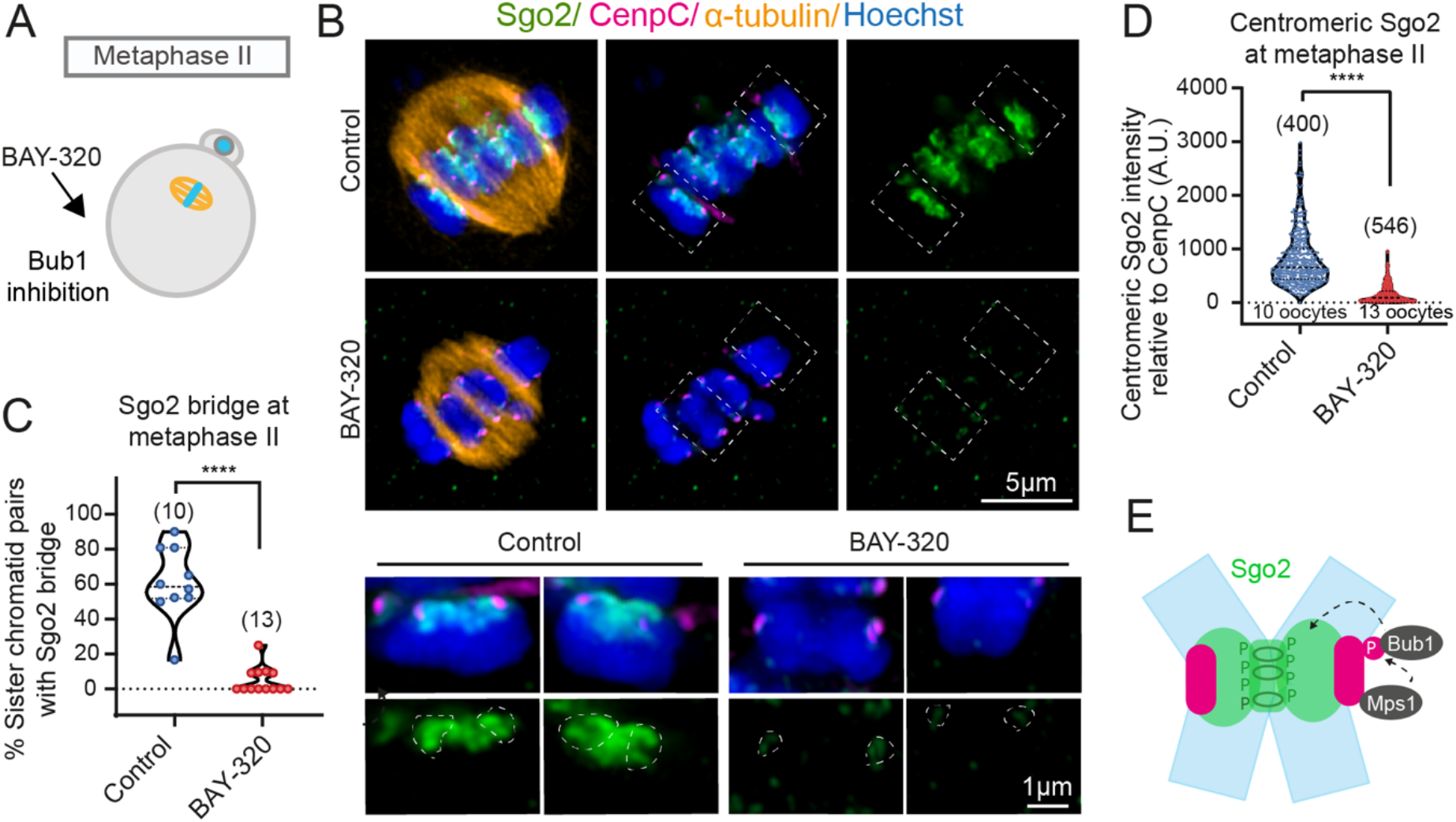
Dependence of Sgo2 localization on Bub1 activity. Loss of Sgo2 localization upon inhibition of Bub1 in metaphase II oocytes. (A) Scheme of the experiment. Metaphase II oocytes from women aged ≤ 36 years were treated with 10μM Bay-320 (to inhibit Bub1^46^) or DMSO (control) overnight and fixed. (B) Representative images of control and Bay-320-treated metaphase II oocytes after immunostaining with antibodies against Sgo2 (green), CenpC (inner kinetochores, magenta) and a-tubulin (microtubules, orange). White boxes with dashed lines indicate chromosomes that have been further magnified below. (C) The percentage of chromatids per oocyte with Sgo2 localization at the pericentromeric bridge was scored in control and Bay-320-treated metaphase II oocytes (\*\*\*\**P* <0.0001, Mann-Whitney test). (D) The relative intensity of the centromeric pool of Sgo2 in metaphase II oocytes from control and Bay-320-treated oocytes were measured relative to CenpC. (****P <0.0001, Mann-Whitney test).Plots show median (dashed black line), 25th and 75th percentiles (dotted black lines).

### Sgo2 protects pericentromeric cohesion

We hypothesised that loss of Sgo2 from the pericentromeric bridge in aged oocytes leaves residual centromeric cohesion vulnerable to cleavage by separase already in anaphase I. This predicts that artificial removal of Sgo2 in meiosis I in younger oocytes would recapitulate the premature loss of centromeric cohesion observed in older oocytes (Figure 3B). To test this idea, oocytes at prophase I (containing a germinal vesicle nucleus), provided by women aged 35 years or younger and undergoing ICSI treatment, were allowed to progress through meiosis I and II in the presence of Reversine to inhibit Mps1, before fixing at metaphase II (Figure 7A). Women in this younger age group were selected for this experiment since their untreated oocytes are expected to retain Sgo2 on the pericentromeric bridge (Figure 1F). As expected, Mps1 inhibition by Reversine treatment from prophase I led to a loss of Sgo2 signal from centromeres and the bridge at metaphase II, while control oocytes retained Sgo2 in both locations (Figure 7B-D). Chromosome alignment defects and reduced inter-sister kinetochore distance at metaphase II were also observed after Mps1 inhibition from prophase I (Figure S6). However, in contrast to oocytes treated with Reversine only at metaphase II (Figure S4B), Mps1 inhibition from prophase I led to a significant increase in single chromatids in metaphase II, indicating loss of centromeric cohesion already in meiosis I (Figure 7E). Therefore, inhibition of Mps1 from prophase I causes single chromatids in metaphase II, while inhibition only in metaphase II does not (Figures S4B and S5C). This demonstrates that Mps1 inhibition from prophase I must abrogate the protection of pericentromeric cohesion in meiosis I, so that separase cleaves pericentromeric cohesin at the same time as arm cohesin in anaphase I. Taken together with the critical role of Mps1 in localizing Sgo2 to pericentromeres (Figure 5B-D; Figure 7B-D), these data strongly suggest that Mps1 protects pericentromeric cohesin in anaphase I through Sgo2 recruitment. Centromeric Sgo2 signal was comparable on pairs and single chromatids after Mps1 inhibition in meiosis I, suggesting that this Sgo2 pool is not an important factor in causing premature cohesin loss (Figure 7F). Therefore, our findings indicate that Sgo2 in the pericentromeric bridge protects pericentromeric Rec8 at the inter-sister kinetochore junction from cleavage during anaphase I, to ensure that robust sister chromatid cohesion is retained at centromeres until metaphase II (Figure 7G). Given that Sgo2 at the bridge is lost in oocytes of older women, our findings indicate that loss of cohesion protection is a likely contributing factor to age-related aneuploidy.

**Figure 7.**
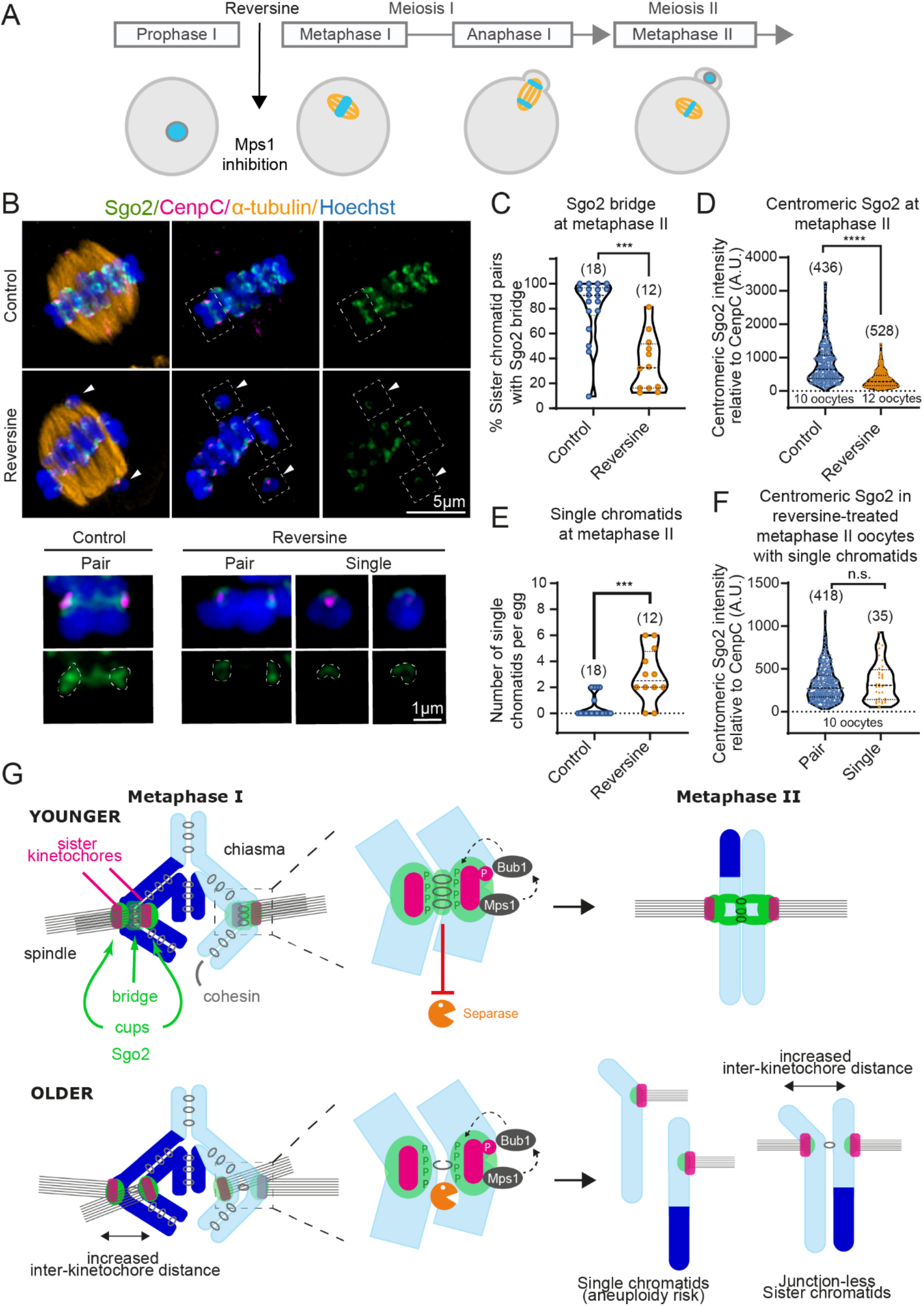
Mps1-dependent Sgo2 protects pericentromeric cohesion during human meiosis. The impact of MPS inhibition on protection of sister chromatid cohesion. (A) Schematic of experiment. Oocytes from women undergoing ICSI aged ≤ 36 years provided at or prior to GVBD were treated with 500nM reversine or DMSO, allowed to mature to metaphase II for up 24 h and fixed. (B) Representative images of oocytes treated with reversine from GVBD stage alongside control oocytes after immunostaining with antibodies against Sgo2 (green), CenpC (inner kinetochore, magenta), a-tubulin (microtubules, orange) and counter staining with Hoechst (chromosomes, blue). White boxes with dashed lines represent chromosomes that have been further magnified below. (C) The percentage of chromatids per oocyte with Sgo2 localization at the pericentromeric bridge was scored (\*\*\**P* = 0.0001; Mann-Whitney test). (D) The relative intensity of the centromeric pool of Sgo2 relative to CenpC. (\*\*\*\**P* <0.0001; Mann-Whitney test). (E) The number of single chromatids observed in metaphase II oocytes after treatment with reversine from GBVD compared to controls (\*\*\**P* <0.0003; Mann-Whitney test). (F) Centromeric Sgo2 was measured in metaphase II oocytes that had single chromatids after reversine treatment from GVBD stage onwards, and values for paired and single chromatids were compared. Sgo2 intensity was measured in arbitrary units relative to CenpC. Plots show median (dashed black line), 25th and 75th percentiles (dotted black lines) (*P* =0.57; Mann-Whitney test). (G) Model for age-dependent loss of cohesin protection. In oocytes from younger women, cohesion is maintained between sister chromatids. Mps1 at the kinetochore recruits Bub1 which phosphorylates histone H2A Thr120 in the centromeric chromatin. This phosphorylation allows for the recruitment of Sgo2 at the centromere and the pericentromeric bridge coincident with cohesin. During anaphase I, Sgo2 protects cohesin within the pericentromeric bridge from separase activity to ensures that sister chromatids remain together at metaphase II and disjoin accurately only upon fertilisation-triggered anaphase II. In oocytes from older women, pericentromeric cohesin deteriorates. This both increases inter-sister kinetochore distances and results in loss of Sgo2 from the pericentromeric bridge. In the absence of Sgo2 at the bridge, residual pericentromeric cohesin is vulnerable to separase-dependent cleavage in anaphase I. Consequently, sister chromatids risk premature separation and such single chromatids at metaphase II which will disjoin randomly at fertilisation resulting in increased incidences of aneuploid conceptions in older women.

## Discussion

A major limitation in the reproductive lifespan of women is the high rates of aneuploidy caused by meiotic errors in the oocyte^1^, which contribute to increasing rates of infertility, miscarriage and chromosomal disorders at older ages. A key driver of this aneuploidy is premature loss of centromeric cohesion^2,7–9^, which prevents sister chromatids aligning on the meiosis II spindle, leading to their mis-segregation in the second meiotic division and production of an aneuploid zygote upon fertilisation. Here we have provided evidence for age-dependent loss of the protector of pericentromeric cohesin as a mechanism that predisposes sister chromatids to increased meiosis II mis-segregation in oocytes from older women.

### Specific loss of Sgo2 from the bridge with age

Our high-resolution analysis of Sgo2 localization in aged human oocytes revealed specific loss of Sgo2 from the bridge – the inter-sister kinetochore junction between sister kinetochores in oocytes. In metaphase II chromosome spreads from younger women, the Sgo2 bridge co-localizes with Rec8 and PP2A, and therefore corresponds to the domain of protected pericentromeric cohesin. A reduction or absence of Sgo2 in this region, as observed in aged oocytes, would render pericentromeric cohesin unprotected and susceptible to cleavage by separase during anaphase I. This premature loss of sister chromatid cohesion is observed as single chromatids in metaphase II (Figure 7G).

We found that Sgo2 loss from the pericentromeric bridge correlated with increased sister kinetochore distance even in oocytes from young women. In mitosis, Sgo1, which protects centromeric cohesin not from separase-dependent cleavage but from non-proteolytic removal by the prophase pathway^47^, is localized to the inner centromere by binding cohesin, while a separate pool of Sgo1 at centromeres is dependent on phosphorylation of H2A by Bub1^32^. If is not clear if Sgo2 is similarly localized through binding cohesin in human oocytes, but these observations raise the possibility that loss of Sgo2 from the bridge is a consequence of the deterioration of cohesin in aged oocytes. In this case, loss of bridge Sgo2 would add a further vulnerability to aged oocytes by exposing the already critically low levels of pericentromeric cohesin to separase-dependent proteolysis. Single chromatids were only rarely observed in metaphase I (Figure S3B), indicating that even in aged oocytes, sufficient cohesin typically remains to hold sister chromatids together. It is only after separase activation at anaphase I that single chromatids are observed (Figure 3B). This indicates that either the age-dependent demise of cohesin leaves insufficient linkages at centromeres to sustain sister chromatid cohesion in the absence of arm cohesin, or that loss of the Sgo2 protector leads to premature cleavage of any centromeric cohesin that survived the age-dependent deterioration. Our demonstration of the importance of Sgo2 in centromeric cohesin maintenance after metaphase I validates the latter possibility. Removal of Sgo2 in prophase I by inhibiting Mps1 led to loss of centromeric cohesion and single chromatids at metaphase II (Figure 7E). These observations also confirm that Sgo2 is the relevant shugoshin for cohesin protection in oocytes and provide a molecular explanation for infertility in a patient with a frameshift *SGO2* mutation^38^.

### Mps1 and Bub1-dependent Sgo2 localization

We found that the kinase activities of both Mps1 and Bub1 are required for localization of human Sgo2 to the centromeric cups and pericentromeric bridge in oocytes. Since Mps1 activity is required for Bub1 localization to kinetochores in human mitotic cells^29,30^ and oocytes^7^ (Figure 5E and F), the most straightforward explanation for these observations is that Mps1 promotes Sgo2 recruitment through localizing Bub1. Bub1 is known to phosphorylate histone H2A on T120 in the centromere to provide a mark that is bound directly by Sgo1 in mitotic cells^32,48^. If a similar mechanism operates in human oocytes, Bub1-dependent H2A-T120 phosphorylation could explain Sgo2 localization at the cups surrounding centromeres. An unresolved issue is how Bub1, which is localized at kinetochores, localizes Sgo2 at the pericentromeric bridge. In mitotic cells, transcription is thought to aid the translocation of centromeric Sgo1 bound to H2A-T120-P to the pericentromere/inner centromere where it binds cohesin^49^ and perhaps Sgo2 re-locates onto pericentromeric cohesin in the bridge in a similar manner in oocytes. However, in mouse oocytes, although Bub1 kinase activity promotes Sgo2 localization to the inter-sister kinetochore junction, Mps1 promotes cohesin protection independently of Bub1^27^. Therefore, Mps1 may also play a direct role in localizing Sgo2 for cohesin protection in human oocytes. Could loss of Sgo2 from the pericentromeric bridge in aged human oocytes be a consequence of a decline in Mps1 or Bub1 function? Intensity measurements found comparable levels of Bub1 at kinetochores in younger and older human oocytes, arguing that loss of the Sgo2 bridge is not a consequence of reduced Bub1 levels as women age^7^. Furthermore, we found that centromeric Sgo2 levels were comparable in oocytes from younger and older women, despite loss of the Sgo2 bridge in the latter (Figure 1F and G). We therefore favour the idea that Bub1 on the kinetochore phosphorylates H2A-T120 in the vicinity to localize the centromeric pool of Sgo2 which is neither directly involved in cohesin protection nor susceptible to loss with advanced maternal age. Instead, as described above, our findings implicate cohesin loss as the key determinant leading to reduced Sgo2 association with the pericentromeric bridge, resulting in an increased vulnerability of this highly specialized pool of cohesin.

### Age dependent loss of cohesion protection as an aneuploidy driver

It is now well-established that erosion of cohesin is a major cause of age-related aneuploidy in human oocytes, with centromeric cohesion weakening being a key driver^2,8,9^. Our study identifies the loss of Sgo2-dependent protection as a further impediment to the safeguarding of centromeric cohesin with age. Since cohesion protection is required only in anaphase I, after ovulation, supplementation of Sgo2 in human oocytes may offer a promising strategy to lessen the effects of aging on aneuploidy predisposition.

## Acknowledgements

We are especially grateful to the women who dated oocytes, embryologists, research nurses and medical consultants at the Edinburgh Fertility and Reproductive Endocrine Centre and the Centre for Reproductive Medicine, University Hospitals Coventry and Warwickshire NHS Trust. We thank all Edinburgh and Warwick colleagues in the Eggs ‘n Embryos research group and Katja Wassmann for helpful discussions. We gratefully acknowledge the Wellcome Centre Optical Imaging Laboratory (COIL) for microscopy support. This work was funded through a Wellcome Collaborator Award [215625] (BPM, GHP, CEC, GMH, ADM, RAA and ALM), a Wellcome Investigator award to ALM [220780] (ALM, BPM, GHP) and core funding for the Wellcome Centre for Cell Biology [203149] (BPM, GHP, DAK and ALM).

---

**Figure S1.**
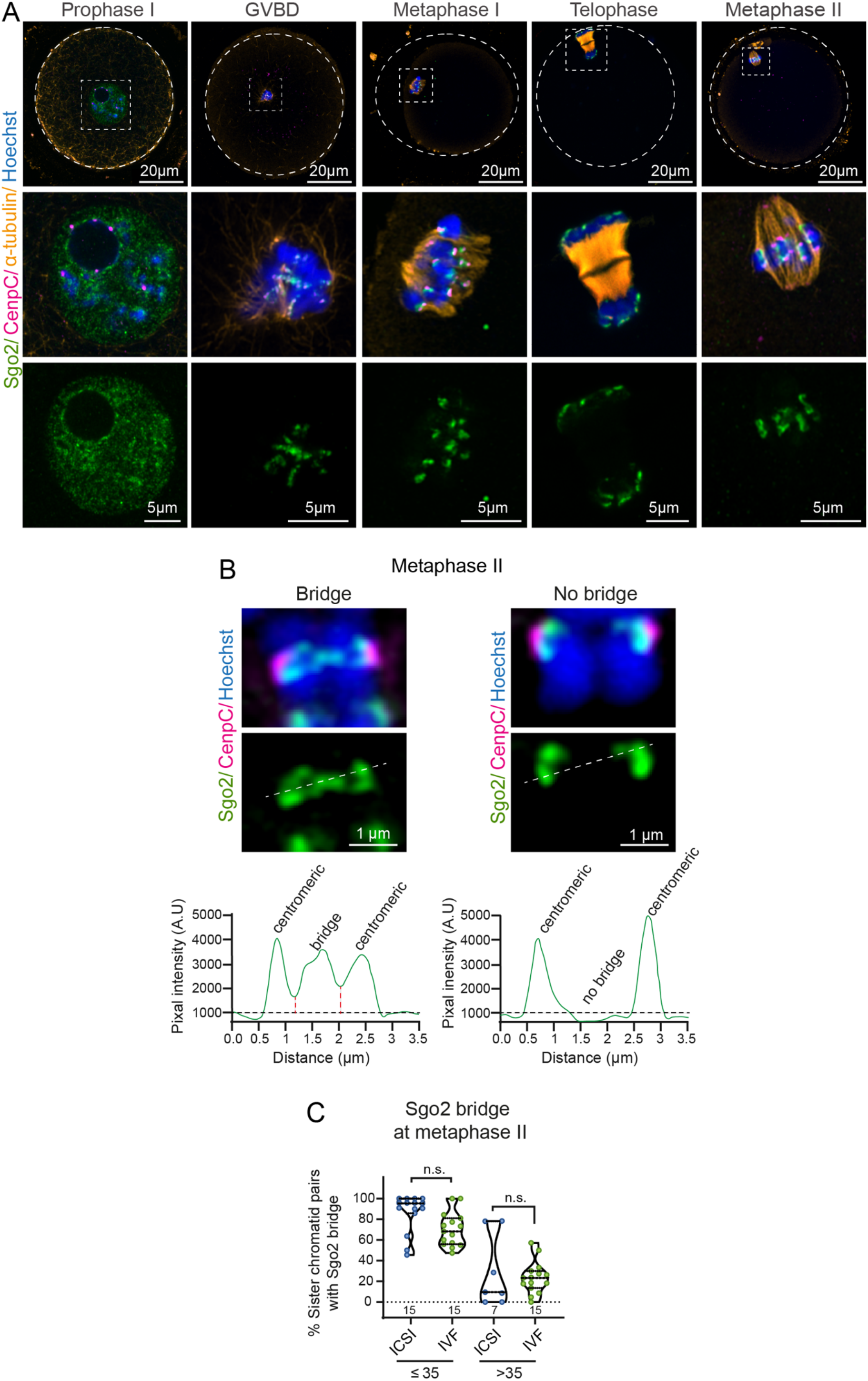
Sgo2 localisation throughout meiosis. Related to Figure 1. (A) Sgo2 localization in GV, GVBD, metaphase I, telophase I, and metaphase II stage oocytes. Oocytes were immunostained with antibodies against Sgo2 (green), CenpC (inner kinetochores, magenta), a-tubulin (microtubules, orange) and counter stained with Hoechst (blue). White circles with dashed lines represent the oocyte circumference. White boxes with dashed lines represent chromosomes masses that have been further magnified below. (B) Representative line scans showing distinction between bridge and centromeric Sgo2 pools. A line was drawn as shown on individual sister chromatid pairs from metaphase II oocytes stained as in (A) and the fluorescence profile of Sgo2 is shown in the graph. (C) Comparison of Sgo2 localization at the pericentromeric bridge in metaphase II oocytes from women stratified by age (≤35 years or >35 years) undergoing ICSI or IVF treatment. Plots show median (dashed black line), 25th and 75th percentiles (dotted black lines). n.s. Not significant (Kruskal-Wallis test).

**Figure S2.**
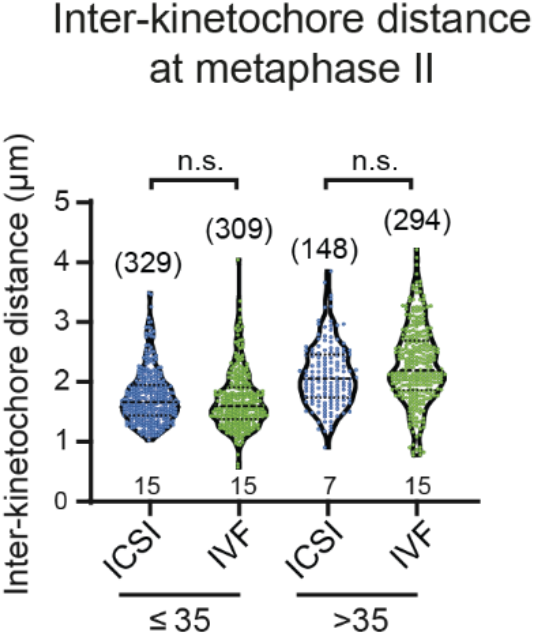
Treatment regime (IVF or ICSI) does not affect age-dependent cohesion loss in metaphase II oocytes. Related to Figure 2. Comparison of inter-kinetochore distance at metaphase II oocytes from women aged ≤35 or >35 years separated into those undergoing ICSI or IVF treatment. Plots show median (dashed black line), 25th and 75th percentiles (dotted black lines). n.s. Not significant (Kruskal-Wallis test).

**Figure S3.**
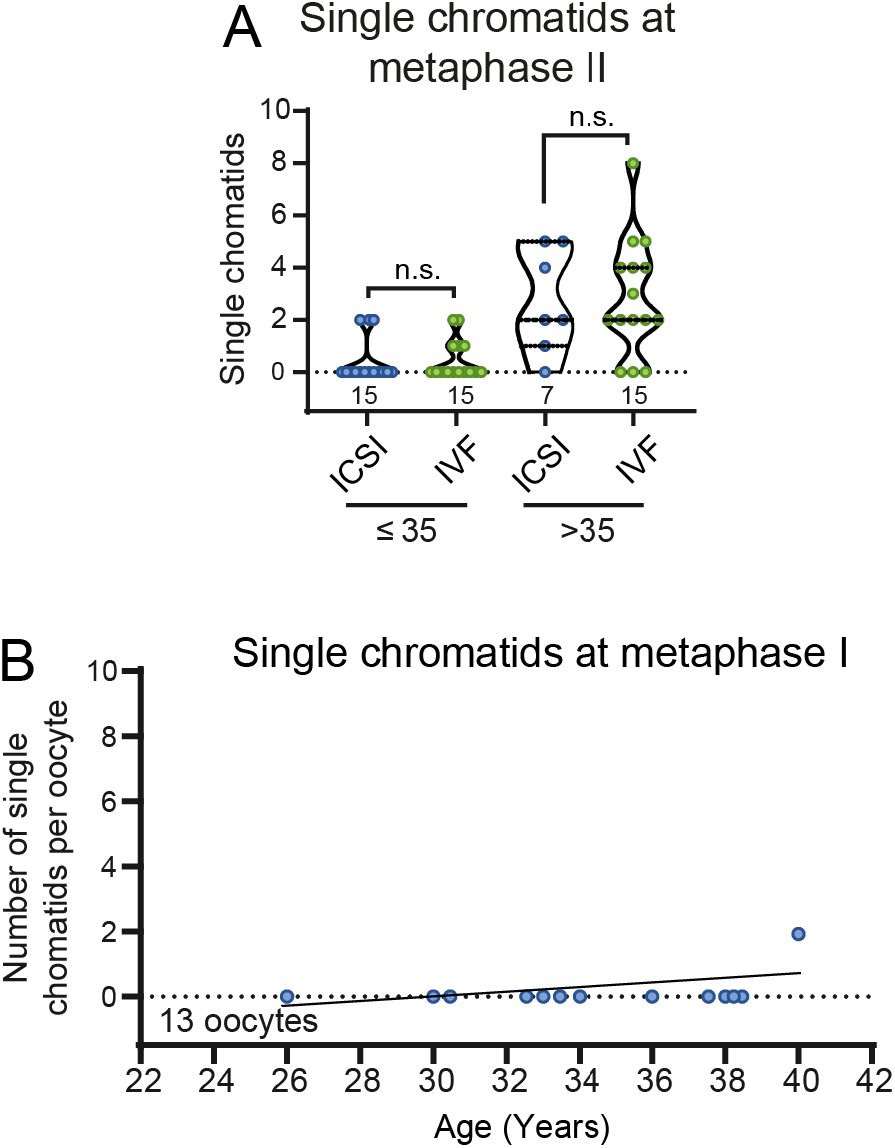
Increased frequency of single chromatids with age is observed only in metaphase II. Related to Figure 3. (A) The number of single chromatids in metaphase I oocytes was scored relative to woman’s age. Metaphase I oocytes were stained as in Figure 1(Kruskal-Wallis test). Plots show median (dashed black line), 25th and 75th percentiles (dotted black lines). *P* values were calculated using the Mann-Whitney test. n.s., Not significant. (B) The number of single chromatids at metaphase I were scored relative to woman’s age (≤35 and >35). Data was fit to a linear regression (R^2^ = 0.1699; *P* = 0.1617).

**Figure S4.**
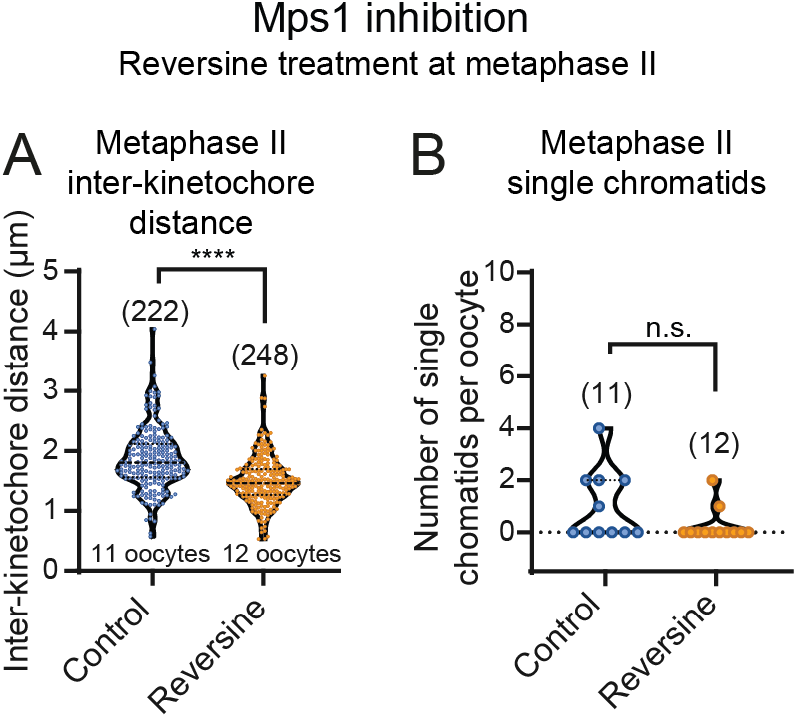
related to Figure 5. Effect of Mps1 inhibition on inter-kinetochore distances and single chromatids. Related to Figure 5. (A) Inter-kinetochore distance and the (B) the number of single chromatids identified in control and metaphase II oocytes from women aged ≤ 36 years treated with reversine as in Figure 5. Plots show median (dashed black line), 25th and 75th percentiles (dotted black lines). ****P <0.0001 (Mann-Whitney test). n.s., Not significant (P=0.12, Mann-Whitney test).

**Figure S5.**
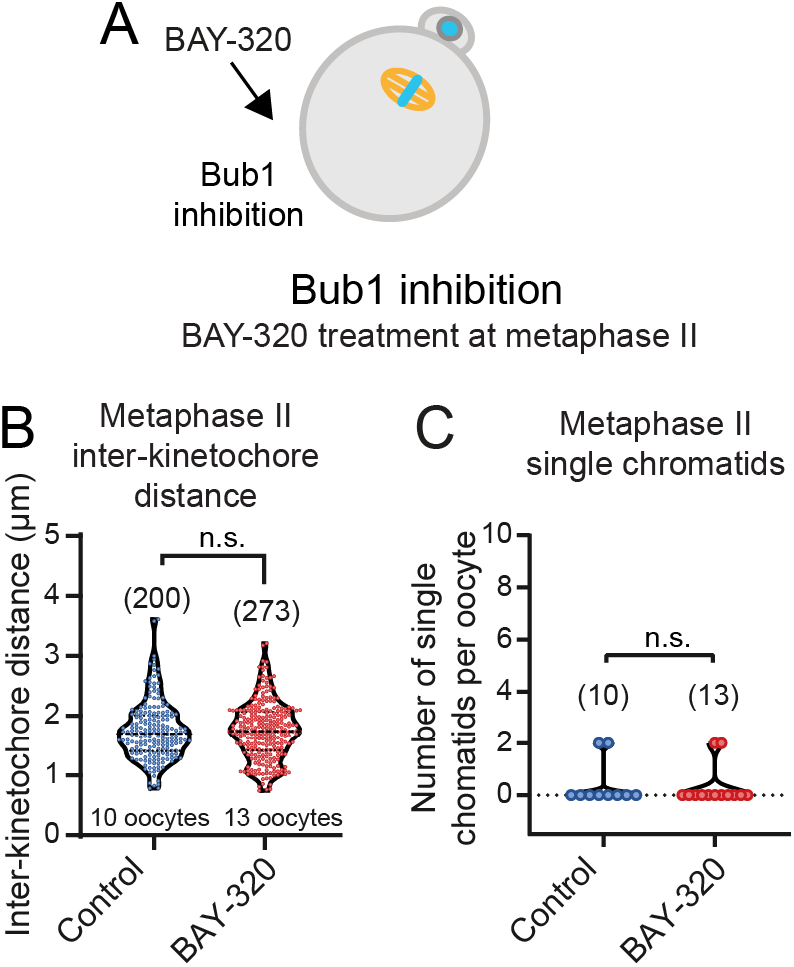
Effect of Bub1 inhibition on inter-kinetochore distances and single chromatids. Related to Figure 6. (A) Inter-kinetochore distance and the (B) the number of single chromatids identified in control and metaphase II oocytes from women aged ≤ 36 years treated with BAY-320 as in Figure 5 Plots show median (dashed black line), 25th and 75th percentiles (dotted black lines). (Mann-Whitney test). n.s. Not significant; Mann-Whitney test (For (A) *P* >0.56; (B) P>0.9999).

**Figure S6.**
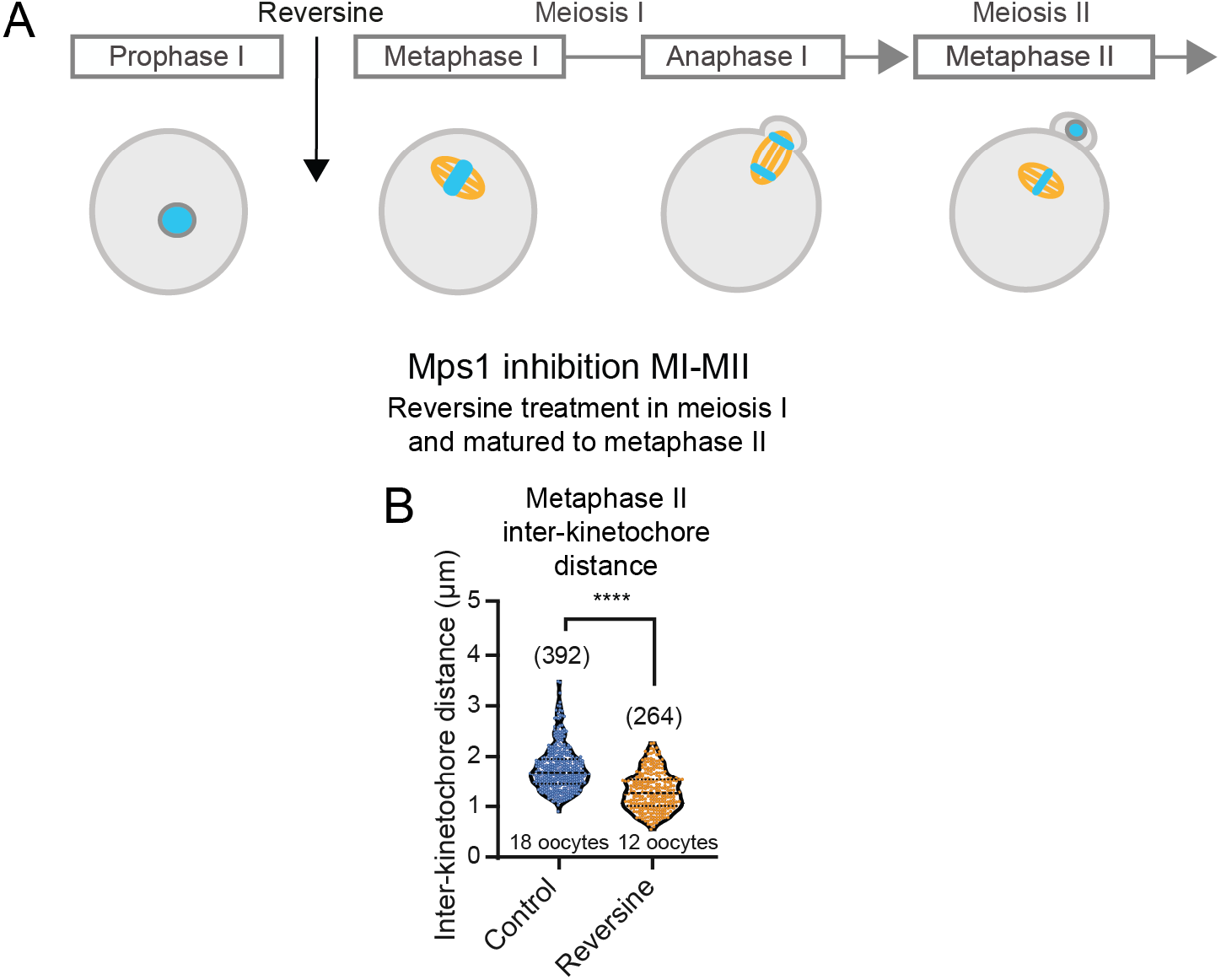
Effect of Mps1 inhibition at GVBD on inter-sister kinetochore distance at metaphase II. Related to Figure 7. (A) Schematic of the experiment reproduced from Figure 7A. (B) Decrease in inter-sister kinetochore distance in oocytes treated with reversine at GVBD and matured to metaphase II as in Figure 7A-D (****P<0.0001, Mann-Whitney test).

## Methods

### Donation of human oocytes to research

The NHS Research Ethics Committee approved the research project (Indicators of Oocyte and Embryo Development, 04/Q2802/26) and all work was conducted under a Research Licence from the Human Fertilisation and Embryology Authority (HFEA; R0155; Indicators of Oocyte and Embryo Development). Informed consent for donation of oocytes and embryos to research was provided by couples undergoing intracytoplasmic sperm injection (ICSI) or in vitro fertilisation (IVF) at the Edinburgh Fertility and Reproductive Endocrine Centre or the Centre for Reproductive Medicine (CRM), University Hospitals Coventry and Warwickshire (UHCW) NHS Trust. Donations were optional and did not affect the treatment received. Couples were aware of the purpose of the research and were not provided compensation. Material donated to research only included oocytes that could not be used for treatment and would have otherwise been disposed of including, immature oocytes and inseminated but unfertilised oocytes. Ovarian stimulation was induced using FSH according to standard clinical protocols, either GnRH agonist or antagonist regimens. Oocytes were cultured in the Vitrolife media suite. After clinical embryologists had completed ICSI or IVF procedures, research material identified and made available for collection by licenced researcher. Immature oocytes arising from ICSI treatments were collected from the clinic between 4-6 hours after oocyte retrieval. Unfertilised or misfertilised metaphase II oocytes from ICSI or IVF were collected from the clinic 3-5 hours after the fertilisation check had been completed by a clinical embryologist ~22 hours after insemination. Cells were transported in G-MOPS PLUS medium (Vitrolife, #10130) at 37°C in a portable incubator (K Systems) for approximately 15 min to the research laboratory. Cells were then cultured in G-IVF PLUS (Vitrolife, #10136) under mineral oil (Merck, # 8042-47-5) at 37°C in 5% O_2_, 6% CO_2_ and 89% N_2_. No oocytes were vitrified or thawed in this study.

A total of 145 oocytes from 91 donors, from November 2019 to December 2022, were used in this study.

### Immunofluorescence

Whole oocytes were fixed and immunostained as previously described with minor modification^9^. Briefly, cells were washed through warmed PHEM buffer (60 mM PIPES, 25 mM HEPES, 10 mM EGTA, 4 mM MgSO4.7H2O; pH 6.9) with 0.25% Triton X-100 at 37°C.

Cells were then fixed with 4% formaldehyde in PHEM buffer with 0.25% Triton X-100 for 30 min and permeabilised further in PBS with 0.25% Triton X-100 for 15 min at room temperature. Cells were then stored in PBS with 0.05% Tween-20 (PBST) at 4°C for up to two weeks until they were immunostained. For immunofluorescence, oocytes were blocked in 3% BSA in PBST at 4°C overnight before being incubated in primary antibodies. Cells were incubated in human polyclonal anti-centromere protein antibody (1:50; Antibodies Incorporated, 15-235), rabbit polyclonal anti-SGOL2 antibody (1:200; Novus Biologicals, NBP1-83567), mouse monoclonal anti-Bub1 antibody (1:100; ThermoFisher Scientific, MA15755), mouse monoclonal antibody against α-tubulin antibody (1:200; T6074, Sigma) and guinea pig polyclonal antibody anti-CenpC antibody (1:200; MBL, PB030) 3% BSA in PBST at 4°C overnight. Following 3 × 20 min washes in 1% BSA in PBST, cells were incubated in goat anti-rabbit Alexa Fluor 488 (1:500; A-11008, ThermoFisher Scientific) goat anti-mouse Alexa Fluor 555 secondary antibody (1:500; A-21422, ThermoFisher Scientific), goat anti-human Alexa Fluor 647 (1:500; A-21445, ThermoFisher Scientific), and goat anti-guinea pig Alexa Fluor 647 secondary antibody (1:500; A-21450, ThermoFisher Scientific) in 3% BSA in PBST for 1 hr at room temperature and then washed 3 × 20 min in 1% BSA in PBST. Oocytes were mounted in ProLon Gold hard-set antifade mountant with DAPI (P36931, ThermoFisher Scientific) on a FluoroDish (FD35-100, WPI) and incubated in the dark overnight at room temperature to allow the mounting medium to harden. Cells were stored in the dark at 4°C and imaged within two weeks of being mounted.

### Chromosome spreads

Chromosomes spreads of human oocytes were prepared by adapting a previously described method for mouse oocytes^50^. Briefly, oocytes were incubated in Tyrode′s Solution, Acidic (T1788, Merck) at 37°C for approximately 20-30 seconds until the zona pellucida dissolved. The oocytes were then washed through three wells containing G-IVF PLUS (10136, Vitrolife) at 37°C. Oocytes were then dropped into a well of a 12 well slide (Epredia, X1XER302W) containing 30μl of fixing solution (3mM DTT, 1% formaldehyde and 0.15% Triton-X-100; PH 9.2-9.3) and allowed to burst. Slides were slowly air dried in slightly open humidified chamber overnight at room temperature. Chromosome spreads were immediately used for immunostaining or frozen at −20°C and used within two weeks. For immunostaining, chromosome spreads were washed for 3 × 5 min in PBST before being incubated in the primary antibodies. Spreads were incubated in mouse monoclonal anti-PP2A Catalytic α antibody (1:50; BD Biosciences, 610556), rabbit polyclonal anti-SGOL2 antibody (1:200; Novus Biologicals, NBP1-83567), mouse polyclonal anti Rec8 antibody (1:50; Novus Biologicals, H00009985-B01P) human anti-CREST antibody/ antisera (1:50 Antibodies Incorporated, 15-235) and guinea pig polyclonal antibody anti-CenpC antibody (1:200; MBL, PB030) diluted in 3% BSA, 7% FBS in PBST. Chromosome spreads were then washed through 1% BSA in PBST and incubated in goat anti-rabbit Alexa Fluor 488 (1:500; A-11008, ThermoFisher Scientific), goat anti-mouse Alexa Fluor 555 secondary antibody (1:500; A-21422, ThermoFisher Scientific) and goat anti-guinea pig Alexa Fluor 647 secondary antibody (1:500; A-21450, ThermoFisher Scientific) in 3% BSA in PBST for 1 hr at room temperature. Chromosome spreads were counterstained with Hoechst 33258 (20 μg/ml) diluted in PBST for 5 min at room temperature and were then mounted using SlowFade Glass Soft-set antifade mountant (ThermoFisher Scientific, S36917) and 1.5H coverslips (VWR, 631-0121). Slides were then sealed with clear nail polish (Revlon) and incubated in the dark overnight at room temperature to allow the mounting medium to set. Spreads were stored in the dark at 4°C and imaged within two weeks of being mounted.

### Human oocyte culture and treatments

Oocytes were cultured in G-IVF PLUS (Vitrolife, #10136) under mineral oil (Merck, # 8042-47-5) at 37°C in 5% O_2_, 6% CO_2_ and 89% N_2_. The stage of oocyte meiosis was visually determined as Prophase I by the presence of a germinal vesicle nucleus (GV) and Metaphase II by the presence of a polar body. Oocytes were matured to prometaphase I, metaphase I and telophase I 10, 12, and 14 hours after nuclear envelope breakdown^27,51^. Oocyte stages were verified by chromosome conformation and tubulin immunolocalisation.

For Mps1 and Bub1 inhibition at metaphase II, oocytes were treated with 500nM reversine (Cayman Chemical Research, 10004412)^27^ or 10μM BAY-320 (MedChemExpress, HY-104000)^46,52^, respectively for 16 hrs prior to fixation, or with an appropriate volume of DMSO as a control. To inhibit Mps1 from prophase I, oocytes were allowed to undergo NEBD and were then incubated with 500nM reversine or 5μM BAY-320, for 20 hours to allow maturation to metaphase II.

### Super resolution imaging

Fixed cells and spreads were imaged using LSM880 or LSM980 laser scanning confocal equipped with an Airyscan 1 or Airyscan 2 detector, respectively (Zeiss UK, Cambridge), using a Plan-APO (63x/1.4 NA) oil objective (Zeiss). 0.14-0.25 μm optical section spacing was used to encompass entire chromatin structures. 405nm, 488nm, 561nm and 639nm lasers were used to detect DAPI and Hoechst staining and Alexa Fluor 488 Alexa Fluor 555 and Alexa Fluor 647, respectively. Emission filters were used to prevent emissions from neighbouring wavelength. Material used for signal intensity measurements were imaged using the same parameters. Airyscan images were subject to 3D Airyscan Processing using the Auto Filter and Medium Strength functions using Zeiss Zen 3.3 (blue edition) software for pixel reassignment and deconvolution to generate 120 nm resolution images. Representative images were prepared using Fiji software (National Institutes of Health).

### Image processing and data analysis

Image processing and quantification were performed with Fiji software. The integrated fluorescence intensity at the centromeres was measured manually using the free hand selection tool and the background fluorescence was measured at four locations on the image and averaged. For determination of fluorescence intensity, the normalized fluorescence (NF), was used as described in the following equation: NF = integrated fluorescence intensity − (area × average background fluorescence). All intensity measurements were normalised to CREST or CenpC signal on the same kinetochore. To measure inter kinetochore distance, kinetochores from sister chromatids were manually identified and distance measurements were taken from their peak intensities. Inter kinetochore distances were measured in 3D using a macro developed by Dr David Kelly at the Wellcome Centre for Cell Biology. All measurement analysis included between 80-100% of available centromere intensities and kinetochores pairs to minimise bias. Chromosomes measured were clearly paired. Ambiguous pairs were eluded from analysis. Sgo2, PP2A and Rec8 between sister chromatids were scored as joined (bridge) or separated (no bridge) manually using Fiji based on an absence of signal between kinetochores, relative to the background. Single chromatids were counted manually in Fiji and were confirmed using SyGlass virtual reality software.

### Statistical analysis and data presentation

Statistical analysis and graphs were generated using Graphpad Prism 9 software (San Diego) and were assembled using Adobe Illustrator. Statistical tests were performed using Mann-Whitney test or Kruskal-Wallis test for non-paired and non-normally distributed data. Chi square test was used to test the association between categorical variables in Figure 4. Linear regression and Sigmodal, 4PL fit were selected based on best fit (R=1), as detailed in the figure legends. P values are designated as *P < 0.05, **P < 0.01, ***P < 0.001, and ****P < 0.0001. Non-significant values are indicated as n.s. The number of replicates are indicated in the figure legends. All violin plots show median (dashed black line), 25th and 75th percentiles (dotted black lines).

## Data availability

All relevant data, Fiji macros and protocols are available from the authors.

## References

1. Hassold, T., and Hunt, P. (2001). To err (meiotically) is human: the genesis of human aneuploidy. Nat Rev Genet 2, 280–291.

2. Gruhn, J.R., Zielinska, A.P., Shukla, V., Blanshard, R., Capalbo, A., Cimadomo, D., Nikiforov, D., Chan, A.C.H., Newnham, L.J., Vogel, I., et al. (2019). Chromosome errors in human eggs shape natural fertility over reproductive life span. Science 365, 1466–1469. 10.1126/science.aav7321.

3. Duro, E., and Marston, A.L.A.L. (2015). From equator to pole: splitting chromosomes in mitosis and meiosis. Genes & development 29, 109–122. 10.1101/gad.255554.114.

4. Tachibana-Konwalski, K., Godwin, J., Van Der Weyden, L., Champion, L., Kudo, N.R., Adams, D.J., and Nasmyth, K. (2010). Rec8-containing cohesin maintains bivalents without turnover during the growing phase of mouse oocytes. Genes and Development 24, 2505–2516. 10.1101/gad.605910.

5. Burkhardt, S., Borsos, M., Szydlowska, A., Godwin, J., Williams, S.A., Cohen, P.E., Hirota, T., Saitou, M., and Tachibana-Konwalski, K. (2016). Chromosome Cohesion Established by Rec8-Cohesin in Fetal Oocytes Is Maintained without Detectable Turnover in Oocytes Arrested for Months in Mice. Current biology : CB 26, 678–685.

6. Duncan, F.E., Hornick, J.E., Lampson, M.A., Schultz, R.M., Shea, L.D., and Woodruff, T.K. (2012). Chromosome cohesion decreases in human eggs with advanced maternal age. Aging cell 11, 1121–1124.

7. Lagirand-Cantaloube, J., Ciabrini, C., Charrasse, S., Ferrieres, A., Castro, A., Anahory, T., and Lorca, T. (2017). Loss of Centromere Cohesion in Aneuploid Human Oocytes Correlates with Decreased Kinetochore Localization of the Sac Proteins Bub1 and Bubr1. Scientific reports 7, 44001.

8. Zielinska, A.P., Holubcová, Z., Blayney, M., Elder, K., and Schuh, M. (2015). Sister kinetochore splitting and precocious disintegration of bivalents could explain the maternal age effect. eLife 4, e11389.

9. Patel, J., Tan, S.L., Hartshorne, G.M., and McAinsh, A.D. (2015). Unique geometry of sister kinetochores in human oocytes during meiosis I may explain maternal age-associated increases in chromosomal abnormalities. Biology open 5, 178–184.

10. Buonomo, S.B., Clyne, R.K., Fuchs, J., Loidl, J., Uhlmann, F., and Nasmyth, K. (2000). Disjunction of homologous chromosomes in meiosis I depends on proteolytic cleavage of the meiotic cohesin Rec8 by separin. Cell 103, 387–398.

11. Klein, F., Mahr, P., Galova, M., Buonomo, S.B., Michaelis, C., Nairz, K., and Nasmyth, K. (1999). A central role for cohesins in sister chromatid cohesion, formation of axial elements, and recombination during yeast meiosis. Cell 98, 91–103.

12. Ferrandiz, N., Barroso, C., Telecan, O., Shao, N., Kim, H.M., Testori, S., Faull, P., Cutillas, P., Snijders, A.P., Colaiácovo, M.P., et al. (2018). Spatiotemporal regulation of Aurora B recruitment ensures release of cohesion during C. elegans oocyte meiosis. Nature communications 9. 10.1038/S41467-018-03229-5.

13. Le, A.H., Mastro, T.L., and Forsburg, S.L. (2013). The C-terminus of S. pombe DDK subunit Dfp1 is required for meiosis-specific transcription and cohesin cleavage. Biology Open 2, 728–738. 10.1242/BIO.20135173/-/DC1.

14. Ishiguro, T., Tanaka, K., Sakuno, T., and Watanabe, Y. (2010). Shugoshin-PP2A counteracts casein-kinase-1-dependent cleavage of Rec8 by separase. Nature cell biology 12, 500–506. 10.1038/NCB2052.

15. Rumpf, C., Cipak, L., Dudas, A., Benko, Z., Pozgajova, M., Riedel, C.G., Ammerer, G., Mechtler, K., and Gregan, J. (2010). Casein kinase 1 is required for efficient removal of Rec8 during meiosis I. Cell cycle (Georgetown, Tex.) 9, 2657–2662. 10.4161/CC.9.13.12146.

16. Brar, G.A., Kiburz, B.M., Zhang, Y., Kim, J.E., White, F., and Amon, A. (2006). Rec8 phosphorylation and recombination promote the step-wise loss of cohesins in meiosis. Nature 441, 532–536.

17. Katis, V.L., Lipp, J.J., Imre, R., Bogdanova, A., Okaz, E., Habermann, B., Mechtler, K., Nasmyth, K., and Zachariae, W. (2010). Rec8 phosphorylation by casein kinase 1 and Cdc7-Dbf4 kinase regulates cohesin cleavage by separase during meiosis. Developmental cell 18, 397–409.

18. Nikalayevich, E., El Jailani, S., Dupré, A., Cladière, D., Gryaznova, Y., Fosse, C., Buffin, E., Touati, S.A., and Wassmann, K. (2022). Aurora B/C-dependent phosphorylation promotes Rec8 cleavage in mammalian oocytes. Current biology : CB 32, 2281-2290.e4. 10.1016/J.CUB.2022.03.041.

19. Riedel, C.G., Katis, V.L., Katou, Y., Mori, S., Itoh, T., Helmhart, W., Galova, M., Petronczki, M., Gregan, J., Cetin, B., et al. (2006). Protein phosphatase 2A protects centromeric sister chromatid cohesion during meiosis I. Nature 441, 53–61.

20. Kitajima, T.S., Sakuno, T., Ishiguro, K., Iemura, S., Natsume, T., Kawashima, S.A., and Watanabe, Y. (2006). Shugoshin collaborates with protein phosphatase 2A to protect cohesin. Nature 441, 46–52.

21. Lee, J., Kitajima, T.S., Tanno, Y., Yoshida, K., Morita, T., Miyano, T., Miyake, M., and Watanabe, Y. (2008). Unified mode of centromeric protection by shugoshin in mammalian oocytes and somatic cells. Nat Cell Biol 10, 42–52.

22. Ogushi, S., Rattani, A., Godwin, J., Metson, J., Schermelleh, L., and Nasmyth, K. (2021). Loss of sister kinetochore co-orientation and peri-centromeric cohesin protection after meiosis I depends on cleavage of centromeric REC8. Developmental cell 56, 3100-3114.e4. 10.1016/J.DEVCEL.2021.10.017.

23. Gryaznova, Y., Keating, L., Touati, S.A., Cladière, D., El Yakoubi, W., Buffin, E., and Wassmann, K. (2021). Kinetochore individualization in meiosis I is required for centromeric cohesin removal in meiosis II. The EMBO journal 40. 10.15252/EMBJ.2020106797.

24. Mengoli, V., Jonak, K., Lyzak, O., Lamb, M., Lister, L.M., Lodge, C., Rojas, J., Zagoriy, I., Herbert, M., and Zachariae, W. (2021). Deprotection of centromeric cohesin at meiosis II requires APC/C activity but not kinetochore tension. The EMBO journal 40. 10.15252/EMBJ.2020106812.

25. Llano, E., Gómez, R.R., Gutiérrez-Caballero, C., Herrán, Y., Sánchez-MartÁn, M., Vázquez-Quiñones, L., Hernández, T., de Alava, E., Cuadrado, A., Barbero, J.L., et al. (2008). Shugoshin-2 is essential for the completion of meiosis but not for mitotic cell division in mice. Genes & development 22, 2400–2413.

26. Rattani, A., Wolna, M., Ploquin, M., Helmhart, W., Morrone, S., Mayer, B., Godwin, J., Xu, W., Stemmann, O., Pendas, A., et al. (2013). Sgol2 provides a regulatory platform that coordinates essential cell cycle processes during meiosis I in oocytes. eLife 2, e01133.

27. El Yakoubi, W., Buffin, E., Cladière, D., Gryaznova, Y., Berenguer, I., Touati, S.A., Gómez, R.R., Suja, J.A., van Deursen, J.M., and Wassmann, K. (2017). Mps1 kinase-dependent Sgo2 centromere localisation mediates cohesin protection in mouse oocyte meiosis I. Nature communications 8, 694.

28. Marston, A.L., and Wassmann, K. (2017). Multiple duties for spindle assembly checkpoint kinases in meiosis. Frontiers in Cell and Developmental Biology 5, 109. 10.3389/fcell.2017.00109.

29. Musacchio, A. (2015). The Molecular Biology of Spindle Assembly Checkpoint Signaling Dynamics. Curr Biol 25, R1002–1018. 10.1016/j.cub.2015.08.051.

30. Sacristan, C., and Kops, G.J.P.L. (2015). Joined at the hip: kinetochores, microtubules, and spindle assembly checkpoint signaling. Trends Cell Biol 25, 21–28. 10.1016/j.tcb.2014.08.006.

31. Kawashima, S.A., Yamagishi, Y., Honda, T., Ishiguro, K.-I., and Watanabe, Y. (2010). Phosphorylation of H2A by Bub1 prevents chromosomal instability through localizing shugoshin. Science (New York, N.Y.) 327, 172–177.

32. Liu, H., Jia, L., and Yu, H. (2013). Phospho-H2A and cohesin specify distinct tension-regulated Sgo1 pools at kinetochores and inner centromeres. Current biology : CB 23, 1927–1933.

33. Ricke, R.M., Jeganathan, K.B., Malureanu, L., Harrison, A.M., and van Deursen, J.M. (2012). Bub1 kinase activity drives error correction and mitotic checkpoint control but not tumor suppression. The Journal of cell biology 199, 931–949.

34. Hodges, C.A., Revenkova, E., Jessberger, R., Hassold, T.J., and Hunt, P.A. (2005). SMC1beta-deficient female mice provide evidence that cohesins are a missing link in age-related nondisjunction. Nat Genet 37, 1351–1355.

35. Lister, L.M., Kouznetsova, A., Hyslop, L.A., Kalleas, D., Pace, S.L., Barel, J.C., Nathan, A., Floros, V., Adelfalk, C., Watanabe, Y., et al. (2010). Age-related meiotic segregation errors in mammalian oocytes are preceded by depletion of cohesin and Sgo2. Current biology : CB 20, 1511–1521.

36. Liu, L., and Keefe, D.L. (2008). Defective cohesin is associated with age-dependent misaligned chromosomes in oocytes. Reproductive biomedicine online 16, 103–112. 10.1016/S1472-6483(10)60562-7.

37. Chiang, T., Duncan, F.E., Schindler, K., Schultz, R.M., and Lampson, M.A. (2010). Evidence that weakened centromere cohesion is a leading cause of age-related aneuploidy in oocytes. Current biology : CB 20, 1522–1528. 10.1016/J.CUB.2010.06.069.

38. Faridi, R., Rehman, A.U., Morell, R.J., Friedman, P.L., Demain, L., Zahra, S., Khan, A.A., Tohlob, D., Assir, M.Z., Beaman, G., et al. (2017). Mutations of SGO2 and CLDN14 collectively cause coincidental Perrault syndrome. Clin Genet 91, 328–332.

39. Chambon, J.-P., Touati, S.A., Berneau, S., Cladière, D., Hebras, C., Groeme, R., McDougall, A., and Wassmann, K. (2013). The PP2A inhibitor I2PP2A is essential for sister chromatid segregation in oocyte meiosis II. Current biology : CB 23, 485–490.

40. Asghar, A., Lajeunesse, A., Dulla, K., Combes, G., Thebault, P., Nigg, E.A., and Elowe, S. (2015). Bub1 autophosphorylation feeds back to regulate kinetochore docking and promote localized substrate phosphorylation. Nat Commun 6, 8364. 10.1038/ncomms9364.

41. Kitajima, T.S., Hauf, S., Ohsugi, M., Yamamoto, T., and Watanabe, Y. (2005). Human Bub1 defines the persistent cohesion site along the mitotic chromosome by affecting Shugoshin localization. Current biology : CB 15, 353–359.

42. Tang, Z., Sun, Y., Harley, S.E., Zou, H., and Yu, H. (2004). Human Bub1 protects centromeric sister-chromatid cohesion through Shugoshin during mitosis. Proc Natl Acad Sci U S A 101, 18012–18017.

43. Santaguida, S., Tighe, A., D’Alise, A.M., Taylor, S.S., and Musacchio, A. (2010). Dissecting the role of MPS1 in chromosome biorientation and the spindle checkpoint through the small molecule inhibitor reversine. The Journal of cell biology 190, 73–87.

44. Maciejowski, J., Drechsler, H., Grundner-Culemann, K., Ballister, E.R., Rodriguez-Rodriguez, J.A., Rodriguez-Bravo, V., Jones, M.J.K., Foley, E., Lampson, M.A., Daub, H., et al. (2017). Mps1 Regulates Kinetochore-Microtubule Attachment Stability via the Ska Complex to Ensure Error-Free Chromosome Segregation. Developmental cell 41, 143-156.e6. 10.1016/J.DEVCEL.2017.03.025.

45. Sliedrecht, T., Zhang, C., Shokat, K.M., and Kops, G.J.P.L. (2010). Chemical Genetic Inhibition of Mps1 in Stable Human Cell Lines Reveals Novel Aspects of Mps1 Function in Mitosis. PLOS ONE 5, e10251. 10.1371/journal.pone.0010251.

46. Baron, A.P., von Schubert, C., Cubizolles, F., Siemeister, G., Hitchcock, M., Mengel, A., Schröder, J., Fernández-Montalván, A., von Nussbaum, F., Mumberg, D., et al. (2016). Probing the catalytic functions of Bub1 kinase using the small molecule inhibitors BAY-320 and BAY-524. eLife 5, e12187.

47. McGuinness, B.E., Hirota, T., Kudo, N.R., Peters, J.M., and Nasmyth, K. (2005). Shugoshin prevents dissociation of cohesin from centromeres during mitosis in vertebrate cells. PLoS Biol 3, e86.

48. Yamagishi, Y., Honda, T., Tanno, Y., and Watanabe, Y. (2010). Two histone marks establish the inner centromere and chromosome bi-orientation. Science (New York, N.Y.) 330, 239–243.

49. Liu, H., Qu, Q., Warrington, R., Rice, A., Cheng, N., and Yu, H. (2015). Mitotic Transcription Installs Sgo1 at Centromeres to Coordinate Chromosome Segregation. Mol Cell 59, 426–436.

50. Chambon, J.-P., Hached, K., and Wassmann, K. (2013). Chromosome spreads with centromere staining in mouse oocytes. Methods Mol Biol 957, 203–212. 10.1007/978-1-62703-191-2_14.

51. Holubcova, Z., Blayney, M., Elder, K., and Schuh, M. (2015). Error-prone chromosome-mediated spindle assembly favors chromosome segregation defects in human oocytes. Science 348, 1143–1147. 10.1126/science.aaa9529.

52. Siemeister, G., Mengel, A., Fernández-Montalván, A.E., Bone, W., Schröder, J., Zitzmann-Kolbe, S., Briem, H., Prechtl, S., Holton, S.J., Mönning, U., et al. (2019). Inhibition of BUB1 Kinase by BAY 1816032 Sensitizes Tumor Cells toward Taxanes, ATR, and PARP Inhibitors In Vitro and In Vivo. Clinical Cancer Research 25, 1404–1414. 10.1158/1078-0432.CCR-18-0628.

